# Defining cellular complexity in human autosomal dominant polycystic kidney disease by multimodal single cell analysis

**DOI:** 10.1101/2021.10.21.465323

**Authors:** Yoshiharu Muto, Eryn E. Dixon, Yasuhiro Yoshimura, Haojia Wu, Kohei Omachi, Andrew J. King, Eric N. Olson, Marvin G. Gunawan, Jay J. Kuo, Jennifer Cox, Jeffrey H. Miner, Stephen L. Seliger, Owen M. Woodward, Paul A. Welling, Terry J. Watnick, Benjamin D. Humphreys

## Abstract

Autosomal dominant polycystic kidney disease (ADPKD) is the leading genetic cause of end stage renal disease and is characterized by the formation and progressive expansion of kidney cysts. Most ADPKD cases arise from mutations in either the *PKD1* or *PKD2* gene but the precise downstream signaling pathways driving cyst growth are not well understood, and relatively few studies investigate human cystic kidney due to sample scarcity. In order to better understand the cell types and states driving human ADPKD progression, we analyzed eight ADPKD and five healthy human kidney samples, generating a single cell multiomic atlas consisting of ~100,000 single nucleus transcriptomes and ~50,000 single nucleus epigenomes. The integrated datasets identified 11 primary cell clusters including most epithelial cell types as well as large endothelial and fibroblast cell clusters. Proximal tubular cells from ADPKD kidneys expressed a failed repair transcriptomic signature characterized by profibrotic and proinflammatory transcripts. We identified the G protein-coupled receptor GPRC5A as specifically upregulated in cyst lining cells derived from collecting duct. The principal cell subpopulation enriched for GPRC5A expression also exhibited increased transcription factor binding motif availability for NF-κB, TEAD, CREB and retinoic acid receptor families and we identified and validated a distal enhancer regulating GPRC5A expression containing these transcription factor binding motifs. This study establishes the single cell transcriptomic and epigenomic landscape of ADPKD, revealing previously unrecognized cellular heterogeneity.

## Introduction

Autosomal dominant polycystic kidney disease (ADPKD) affects approximately 1 in 400 ~1000 individuals worldwide and it is the most common inherited cystic disease^1^. Cyst growth and expansion in ADPKD ultimately destroys normal kidney tissue leading to CKD and often progressing to end stage kidney disease (ESKD). Most cases are characterized by mutations in either the *PKD1* or *PKD2* genes that encode polycystin-1 (PC-1) and polycystin-2 (PC-2), respectively. PC-1 and PC-2 are transmembrane proteins that form a heterodimeric complex localized to the primary cilium, plasma membrane and endoplasmic reticulum. Some evidence suggests that PC-1 and PC-2 sense fluid flow and regulate intracellular calcium levels^2^, but this remains controversial and the primary function of these proteins is undefined. Other pathways that have been implicated in PC-1 and PC-2 signaling include cAMP, mammalian target of rapamycin complex (mTORC), WNT, metabolic pathways including glycolysis and mitochondrial function^1,2^. The vasopressin receptor antagonist tolvaptan has been approved for slowing the progression of ADPKD and acts by decreasing cAMP concentration, although therapy is associated with polyuria which can limit tolerance^3^. Accordingly, the development of new therapeutic approaches to ADPKD is of paramount importance.

Recent advances in single cell or single nucleus RNA sequencing (scRNA-seq or snRNA-seq) technologies have advanced our understanding of cell types and states present in both healthy and diseased human kidney^4,5^. snRNA-seq in particular is well suited to the analysis of human tissue since it is compatible with cryopreserved samples and we have demonstrated comparable sensitivity to scRNA-seq^6^. Recent single cell profiling approaches have been extended to include the epigenome. The single nucleus assay for transposase-accessible chromatin sequencing (snATAC-seq) technique utilizes hyperactive Tn5 transposase to map open chromatin at single cell resolution^7,8^. The resulting large datasets can be used to predict cis-regulatory DNA networks and transcription factor activity^9,10^, providing complementary information to snRNA-seq^11^. We and others have recently reported multimodal single cell atlases of healthy and diseased human or mouse kidneys and leveraged these atlases to redefine cellular heterogeneity, demonstrating the potential utility of multimodal single cell analyses to understand kidney biology^12–14^. Furthermore, recent advances in epigenetic editing technology with dCas9 fusion protein enables us to validate gene regulatory networks predicted by snATACs-eq analysis^15^.

Here, we have performed snRNA-seq and snATAC-seq on eight human ADPKD kidneys to understand the cell states and dynamics in late stage ADPKD at single cell resolution. We successfully identifying previously unrecognized subpopulations and their molecular signatures in cyst lining cells and other cell types. Most proximal tubular cells in ADPKD kidneys had adopted pro-inflammatory, pro-fibrotic failed repair transcriptomic signature. We could also identify proinflammatory fibroblast and endothelial cell subtypes with evidence of intercellular communication networks present only in ADPKD but not healthy kidneys. We observed specific upregulation of the G protein-coupled receptor GPRC5A in large collecting duct cysts, and we identify a distal *GPRC5A* enhancer regulating its expression in ADPKD principal cells. Our study represents the first multimodal single cell atlas of human ADPKD and reveals new cell states associated with late stage disease.

## Results

### Single cell transcriptional and chromatin accessibility profiling on ADPKD kidneys

We performed snRNA-seq on eight ADPKD and five control adult kidney samples with 10X Genomics Chromium Single Cell 3’ v3 chemistry (Fig. 1a). The ADPKD patients ranged in age from 35 to 61 years and included men (n=4) and women (n=4). All patients had ESKD requiring kidney transplantation at the time of sample collection; 1 patient had been on maintenance dialysis for 6 months prior to transplant and other patients (n=7) received pre-emptive transplantation (Supplementary Table 1). Five control kidneys samples were from were from partial nephrectomy samples in patients with preserved renal function (mean sCr = 1.07 mg/dl) that ranged in age from 50 to 62 years and included men (n=3) and women (n=2). These five samples have been reported previously using the 10X Chromium 5’ Chemistry^12^, however we generated new libraries from these samples using the 3’ chemistry in order to allow direct comparison with the ADPKD samples. After batch quality control (QC) filtering and preprocessing, ADPKD or control snRNA-seq datasets were integrated with Seurat^16^ and visualized in UMAP space to annotate cell clusters (Supplementary Fig. 1a, Supplementary Fig. 2a, See also Methods). Interestingly, lineage marker expression was largely preserved in ADPKD kidneys (Supplementary Fig. 2a), allowing the assignment of cell of origin for cyst cells. All major tubular cell types were identified in both ADPKD and control datasets (Supplementary Fig. 1a, Supplementary Fig. 2a) but leukocyte clusters were only detected in ADPKD dataset (Supplementary Fig. 2a). After removing doublets and low-quality clusters, we obtained a total of 102,710 nuclei by snRNA-seq; 62,073 nuclei from ADPKD and 40,637 nuclei from control kidneys (Supplementary Fig. 1b, Supplementary Fig. 2b). Control kidney datasets had more unique genes and transcripts per cell than ADPKD samples, likely reflecting increased RNA degradation during dissociation from the more fragile ADPKD tissue (Supplementary Fig. 3, See also Methods). ADPKD and control kidney datasets were then integrated in Seurat, and batch correction was performed with the R package “Harmony”^17^ (Fig. 1a, b). Differentially expressed genes were identified (Supplementary Data1-3, Fig. 1c), and cell types were assigned to each of the unsupervised clusters based on lineage markers (Fig. 1b, c, Supplementary Table 2). The two smallest clusters expressed uroepithelial marker genes (*UPK3A*, *PSCA*), suggesting that they were probably uroepithelium (Fig. 1b). Nearly all of these cells (~99%) were from one patient (PKD8).

**Figure 1.**
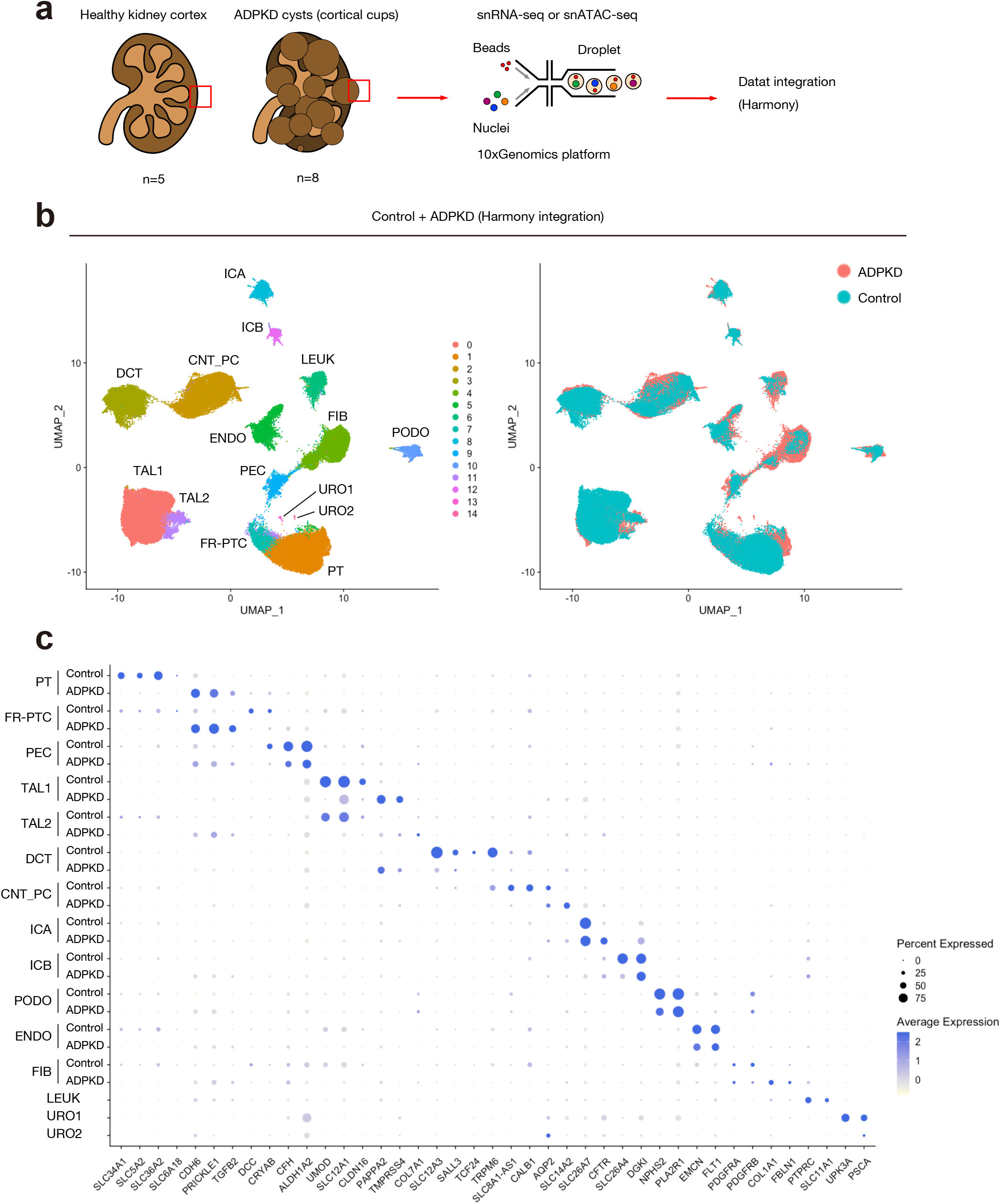
Single nucleus transcriptional profiling on human ADPKD kidneys. (**a**) Graphical abstract of experimental methodology. n=8 human ADPKD kidneys and n=5 control kidneys were analyzed with snRNA-seq and snATAC-seq. Bafailed repair cell states have been identified in snRNA-seq tch effect on the integrated datasets was corrected with Harmony. See Method section for detail. (**b**) UMAP plot of integrated snRNA-seq dataset with annotation by cell type (left) or disease condition (right). PT, proximal tubule; FR-PTC, failed-repair proximal tubular cells; PEC, parietal epithelial cells; TAL, thick ascending limb of Henle’s loop; DCT, distal convoluted tubule; CNT_PC, connecting tubule and principle cells; ICA, Type A intercalated cells; ICB, Type B intercalated cells; PODO, podocytes; ENDO, endothelial cells; FIB, fibroblasts; LEUK, leukocytes; URO, uroepithelium. (**c**) Dot plot of snRNA-seq dataset showing gene expression patterns of cluster-enriched markers for ADPKD or control kidneys. For LEUK and URO1/2 clusters, data from ADPKD kidneys were shown. The diameter of the dot corresponds to the proportion of cells expressing the indicated gene and the density of the dot corresponds to average expression relative to all cell types.

snATAC-seq (10X Genomics Chromium Single Cell ATAC v1) was also performed on the same ADPKD samples to profile single cell chromatin accessibility. snATAC-seq datasets on control kidneys were previously described^12^. Multi-omic integration and label transfer with Seurat was performed on ADPKD and control kidney datasets^11^ (Fig. 2a, See also Methods). The prediction scores for label transfer in the healthy kidneys is higher than that of the ADPKD dataset, likely reflecting the lower gene detection per cell in the ADPKD samples (Supplementary Fig. 4a). The snATAC-seq datasets were filtered using an 80% confidence threshold for cell type assignment to remove heterotypic doublets, and we obtained 33,621 nuclei for control and 17,365 nuclei for ADPKD. Finally, ADPKD and control snATAC-seq datasets were integrated with Harmony (Fig. 2a) and visualized in UMAP space (Fig. 2b), and each cluster was annotated based on gene activities (Fig. 2b). We confirmed that snATAC-seq cell type predictions obtained by label transfer (Supplementary Fig. 4c, d) and curated annotations of unsupervised clusters based on gene activities (Fig. 2b–d, Supplementary Table 3) were largely consistent. We performed downstream analyses with these cell type assignments based on unsupervised clustering and gene activities of lineage markers (Fig.2b-d, Supplementary Data 4-12).

**Figure 2.**
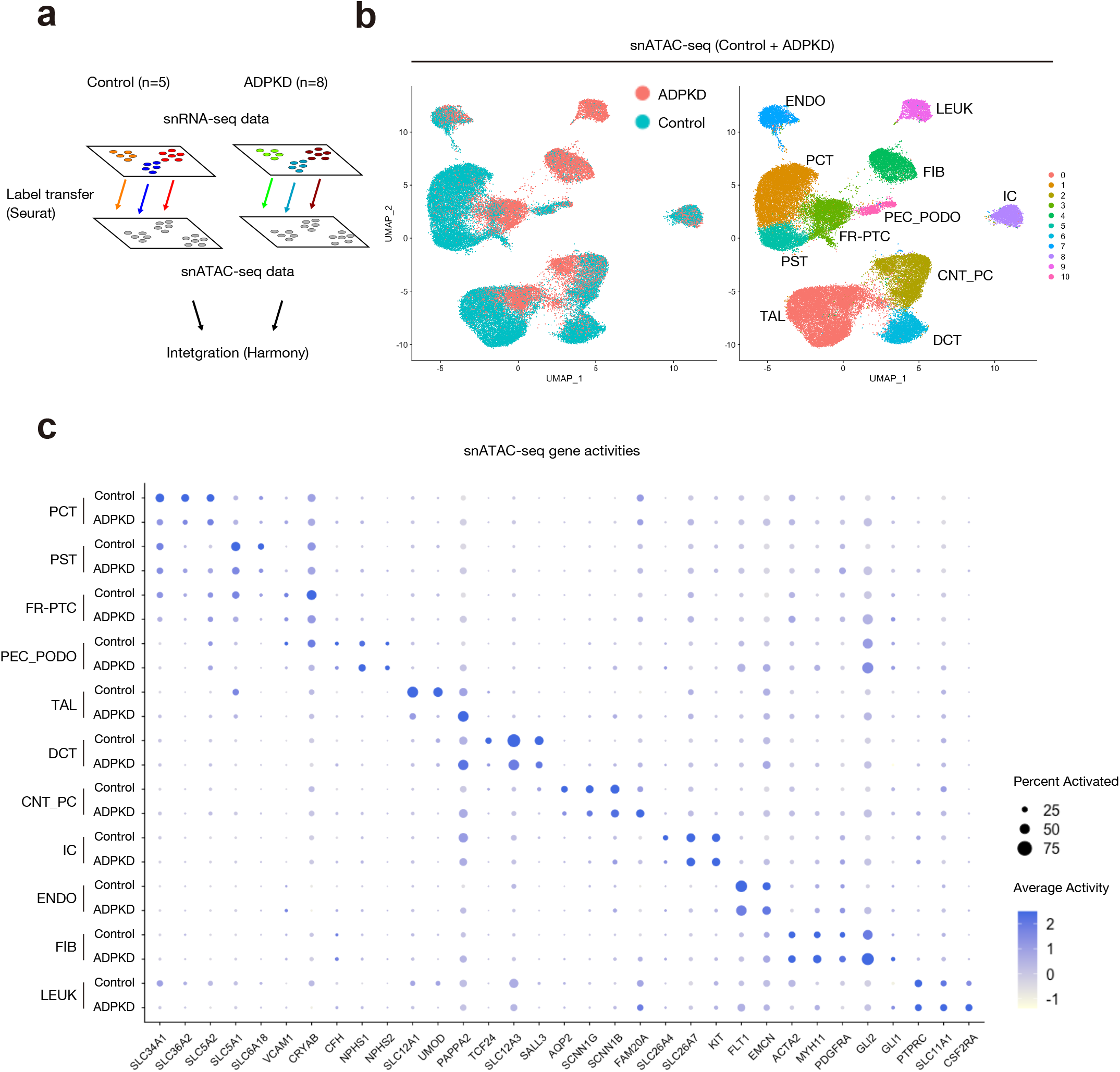
Single nucleus chromatin accessibility profiling on human ADPKD kidneys. (**a**) Graphical abstract of multimodal integration strategy for the snATAC-seq datasets. The integrated ADPKD or control snATAC-seq datasets were label-transferred from cognate snRNA-seq datasets, and the snATAC-seq datasets were filtered using an 80% prediction score threshold for cell type assignment. After filtering, control and ADPKD datasets were merged, and batch effect was corrected with Harmony. See Supplementary Fig. 4 and Method section for detail. (**b**) UMAP plot of snATAC-seq dataset with annotation by disease condition (left) or gene activities-based cell type assignments (right). PCT, proximal convoluted tubule; PST, proximal straight tubule. (**c**) Dot plot of snATAC-seq dataset showing gene activity patterns of cluster-enriched markers for control or ADPKD kidneys. The diameter of the dot corresponds to the proportion of cells with detected activity of indicated gene and the density of the dot corresponds to average gene activity relative to all cell types.

### Proximal tubular cells exhibit a failed repair cell state in ADPKD kidneys

We observed proximal tubular cells (PTC) expressing *VCAM1* in both ADPKD and control kidney datasets (Supplementary Fig. 1a, Supplementary Fig. 2a). *Vcam1*+*Ccl2*+ PTC were recently described as failed-repair proximal tubular cells (FR-PTC) with a proinflammatory and profibrotic transcriptional profile in mouse kidneys ^18–20^. To further characterize the role of FR-PTC in ADPKD progression, we performed unsupervised clustering on PT in our snRNA-seq dataset and identified two subpopulations; N-PTC (<u>Normal</u> proximal tubular cells) and FR-PTC (Fig. 3a, b). We found that most of the PT in ADPKD kidneys were FR-PTC, while most of PT in control were N-PTC (Fig. 3b). The differentially expressed genes in FR-PTC included *VCAM1*, *CCL2* and *HAVCR1* along with *CDH6* (Supplementary Data 13). We previously reported that VCAM1+ cells existed in a scattered fashion (~4%) in PT of healthy subject kidneys^12^. CDH6 is a marker of PT progenitors in developmental kidneys^21^. Further, CDH6 expression has been reported to be up-regulated in renal clear carcinoma^22^ (RCC), arising from VCAM1+CD133+ PTC after kidney injury^23^. Immunostaining of ADPKD samples identified VCAM1+ cells mainly in atrophic tubules and relatively small cysts (Fig. 3c). In contrast, the distal nephron marker CDH1 was identified in large cysts (Fig. 3c).

**Figure 3.**
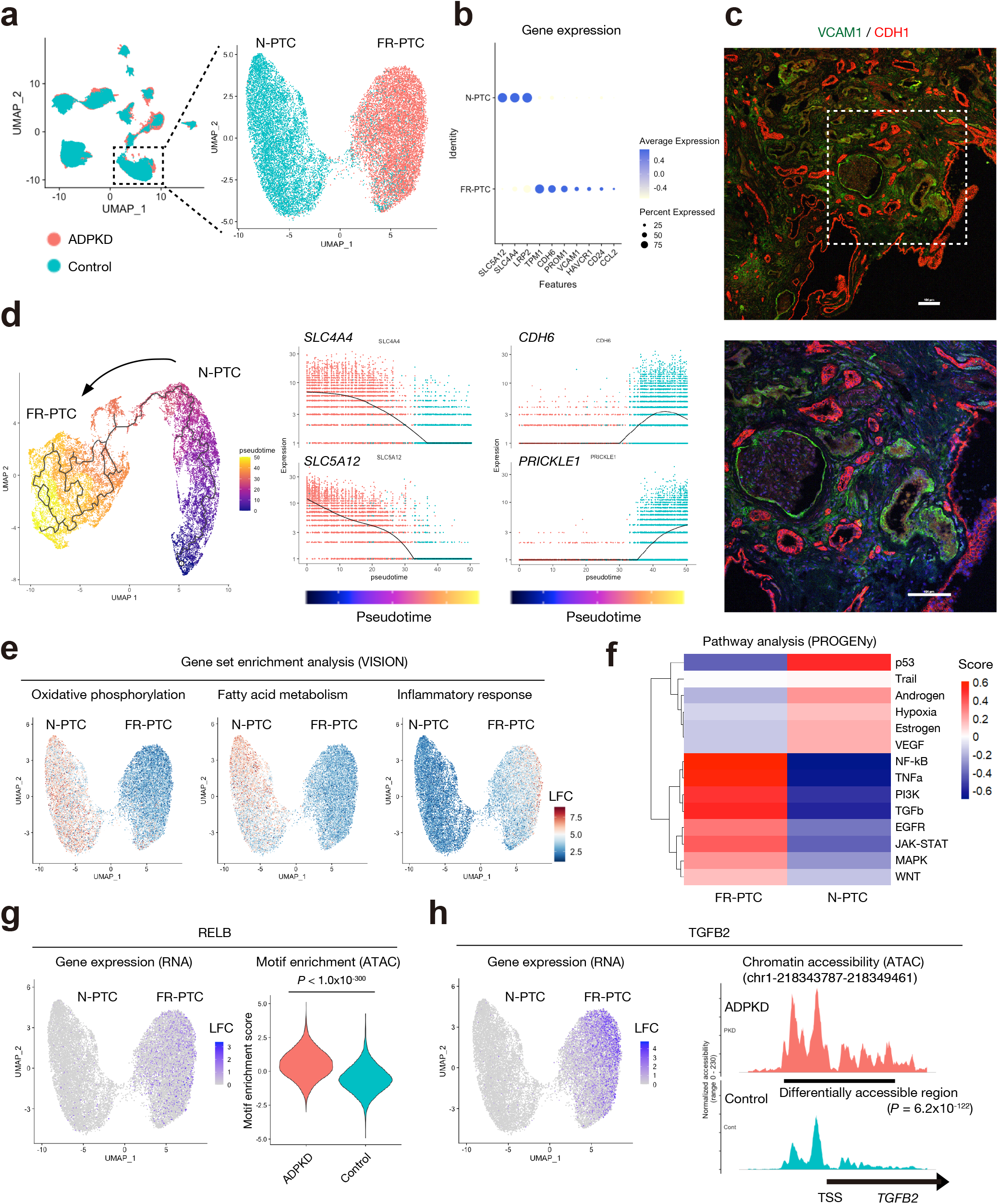
Proximal tubular cells express a failed repair molecular signature in ADPKD kidneys. (**a**) Subclustering of PT on the UMAP plot of snRNA-seq dataset. N-PTC, normal proximal tubular cells; FR-PTC, failed-repair proximal tubular cells. (**b**) Dot plots showing gene expression patterns of the genes enriched in each of PT subpopulations. The diameter of the dot corresponds to the proportion of cells expressing the indicated gene and the density of the dot corresponds to average expression relative to all PT cells. (**c**) Immunofluorescence staining for VCAM1 (green) and CDH1 (red) in the ADPKD kidney sections (representative image of n=3 samples) with 100x (upper) or 200x (lower) field. VCAM1 was expressed in PEC and tubular cells without CDH1 expression. Scale bar indicates 100 μm. (**d**) Pseudotemporal trajectory from N-PTC to FR-PTC using snRNA-seq was generated with Monocle3 (left), and gene expression dynamics along the pseudotemporal trajectory from N-PTC to FR-PTC are shown (right); *SLC4A4* (upper middle), *SLC5A12* (lower middle), *CDH6* (upper right) and *PRICKLE1* (lower right). (**e**) UMAP plot showing enrichment of gene set for oxidative phosphorylation (left), fatty acid metabolism (middle) or inflammatory response (right) at single cell level. The color scale represents a normalized log-fold-change (LFC). (**f**) Heatmap showing pathway enrichment on PCT subpopulations with PROGENy. The color scale represents pathway enrichment score. (**g**) UMAP plot displaying *RELB* gene expression in PT subtypes (left) or violin plot showing RELB motif enrichment score for PT in ADPKD or control dataset. The color scale represents a normalized LFC. (**h**) UMAP plot displaying *TGFB2* gene expression in PT subtypes (left) and fragment coverage (frequency of Tn5 insertion) around the DAR (DAR +/−1 Kb) on the *TGFB2* locus of ADPKD (upper right) or control (lower right) PT. The color scale for each plot represents a normalized LFC. Bonferroni adjusted p-values were used to determine significance.

To characterize cell changes on the failed repair trajectory, we performed pseudotemporal ordering with Monocle^24–26^ from N-PTC to FR-PTC. This revealed gradual reduction in the expression of healthy PT markers and increased expression of FR-PTC marker genes (Fig. 3d). We next analyzed the PT subcluster data using Vision, a tool for annotating the sources of variation in single cell RNA-seq data^27^. FR-PTC lost oxidative phosphorylation gene signatures and gained inflammatory signatures (Fig. 3e), consistent with the previously reported mitochondrial dysfunction in human kidneys with chronic kidney disease (CKD) and inflammatory responses caused by loss of mitochondrial integrity^28^. This was further supported by PROGENy pathway analyses^29^ (Fig. 3f). *Relb* expression has been shown to be up-regulated in mitochondrial dysfunction caused by *Tfam* deletion in a mouse model^28^. In agreement with these findings, *RELB* gene expression was increased in FR-PTC, and the RELB binding motif was enriched in accessible chromatin in ADPKD PT (Fig. 3g, Supplementary Data 14) compared to control PT. Among the most differentially up-regulated genes between FR-PTC and N-PT, we identified *TGFB2* (P value adjusted with Bonferroni correction [*P adj.*] < 1.0 ×10^−300^, Log fold change = 2.22, Fig. 3h), which is known to promote fibrosis of various organs^30^, including the kidneys^31^. The genomic region around the transcription start site (TSS) of *TGFB2* in the ADPKD PT was significantly more accessible compared to that of control kidneys as well (Fig. 3h, Supplementary Data 15), consistent with its gene expression. Interestingly, the expressions of TGFβ receptors were also up-regulated in FR-PTC. Furthermore, the binding motif of SMADs that are downstream effectors of TGFβ signaling was enriched in ADPKD PT in snATAC-seq data (Supplementary Fig. 5). These findings suggest that TGFβ secreted from FR-PTC may be acting in an autocrine or paracrine fashion in addition to signaling to other surrounding cells, including myofibroblasts. Collectively, these findings indicate that the FR-PTC increased in ADPKD kidneys adopt a pro-inflammatory cell state.

### Hedgehog pathway activation among fibroblast subsets in ADPKD kidneys

CKD accompanying ADPKD is associated with interstitial fibrosis^32^. To characterize the alteration in molecular signatures of interstitial cells in ADPKD kidneys, we performed subclustering on fibroblast (FIB) clusters, resulting in separation of 7 subclusters (Fig. 4a,b). FIB1 and FIB2 expressed *PDGFRB,* and they were detected in both ADPKD and control kidneys. In contrast, *ACTA2*+ myofibroblasts (MyoFIB) were exclusively detected in ADPKD kidneys. There was also another ADPKD-specific cluster (PKD-FIB) with a distinct molecular signature (Fig. 4b, Supplementary Data16). PKD-FIB expressed *IL6* and *FGF14* at high levels (Fig. 4b, Supplementary Fig. 6a,b). Two uncharacterized clusters (Unknown1 and Unknown2), expressed tubular cell markers (*SLC12A1*, *SLC34A1* and *LRP2*) that were detected in both ADPKD and control kidneys, most likely residual doublets despite our use of Doubletfinder^33^. Each of these subtypes had variable expression levels of fibroblast marker genes (Fig. 1c, Fig. 4c), suggesting cellular heterogeneity of fibroblasts. Pathway analysis with PROGENy^29^ indicated TGFβ signaling pathway activation in the MyoFIB cluster, as well as TNFα-induced NF-κB activation in the PKD-FIB cluster (Fig. 4d). Since *TGFB2* expression was up-regulated in FR-PTC (Fig. 3h), cyst FR-PTC may be driving interstitial myofibroblast proliferation. *TNF* expression was mainly detected in principal cells of ADPKD kidneys (Supplementary Fig. 6c), suggesting a similar paracrine signaling relationship between collecting duct-derived cysts and interstitial PKD-FIB in ADPKD.

**Figure 4.**
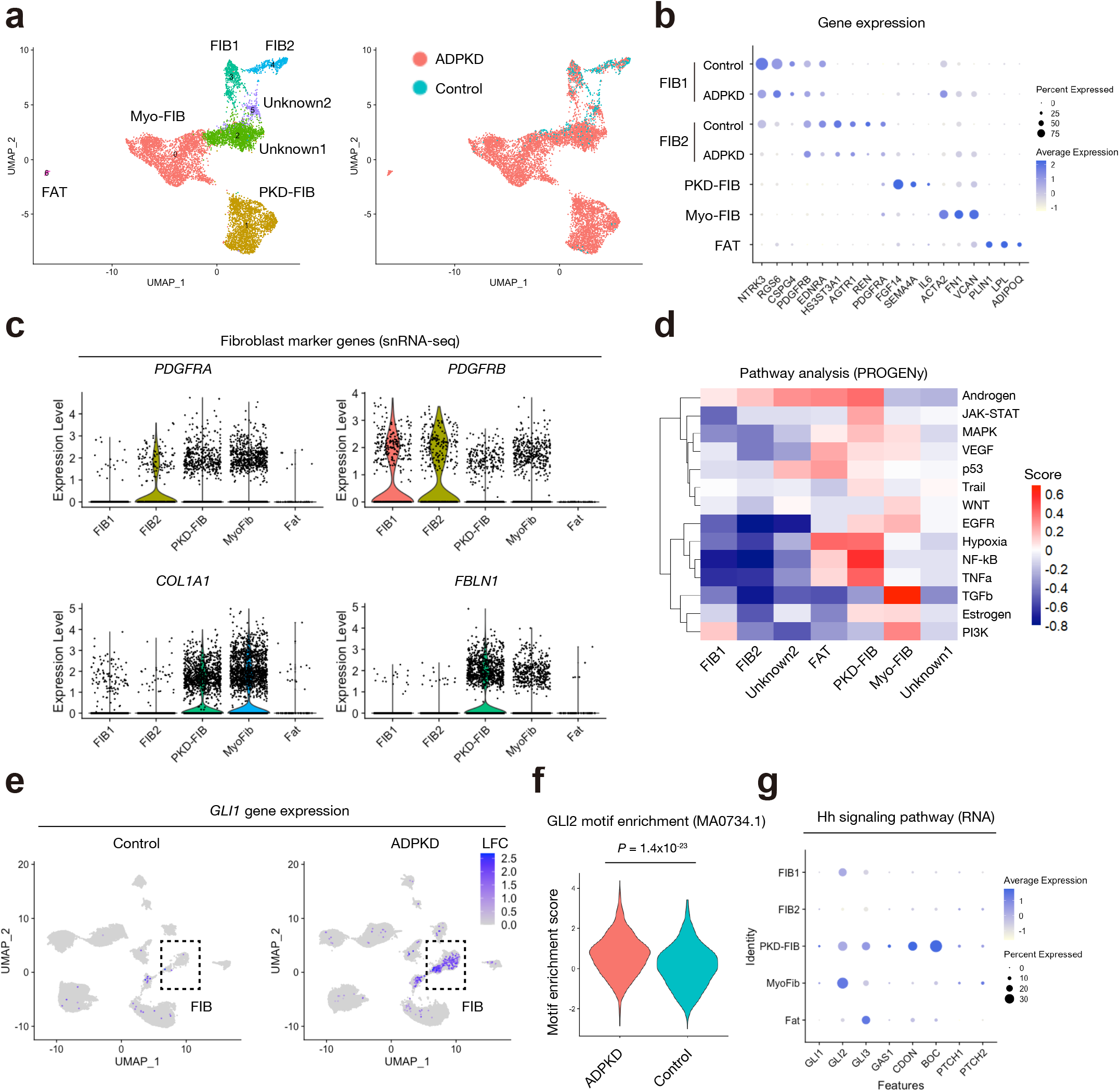
Hedgehog signaling pathway activation among fibroblast subsets in ADPKD kidneys. (**a**) Subclustering of FIB on the UMAP plot of snRNA-seq dataset with annotation by subtype (left) or disease condition (right). MyoFIB, Myofibroblast; PKD-FIB, ADPKD-specific fibroblast subtype; FAT, adipocytes. (**b**) Dot plots showing gene expression patterns of the genes enriched in each of FIB subpopulations. For FIB1 and FIB2, control and ADPKD data were individually shown. The diameter of the dot corresponds to the proportion of cells expressing the indicated gene and the density of the dot corresponds to average expression relative to all FIB cells. (**c**) Violin plot showing fibroblast marker gene expressions among FIB subclusters; *PDGFRA* (upper left), *PDGFRB* (upper right), *COL1A1* (lower left) and *FBLN1* (lower right). (**d**) Heatmap showing pathway enrichment on FIB subpopulations with PROGENy. The color scale represents pathway enrichment score. (**e**) UMAP plot displaying gene expressions of *GLI1* in ADPKD (right) or control kidneys (left). The color scale represents a normalized log-fold-change (LFC). (**f**) Violin plot showing GLI2 motif enrichment score for ADPKD or control kidneys. (**g**) Dot plots showing gene expression patterns of the genes related to hedgehog (Hh) signaling pathway. Bonferroni adjusted p-values were used to determine significance.

Interstitial fibrosis is associated with hedgehog signaling (Hh) pathway activation^34,35^. Consistent with this, the Hh pathway effector and readout of Hh pathway activation *GLI1* was highly expressed in FIB of ADPKD kidneys (Fig. 4e), especially in MyoFIB and PKD-FIB (Supplementary Fig. 6d). *GLI1* gene expression is regulated by GLI2^36^, and GLI2 binding motifs showed enriched accessibility in ADPKD fibroblasts (Fig. 4f, Supplementary Data 12). We also found that the PKD-FIB cluster expressed Hh pathway modulators *GAS1*, *CDON* and *BOC*^37^ (Fig. 4g), suggesting that sensitivity to Hh ligands may differ between MyoFIB and PKD-FIB cluster. Endothelial cells were the primary Hh ligand producing cell type (next section). Together, these findings suggest heterogeneity of fibroblasts with Hh pathway activation in ADPKD kidneys.

### Transcriptional and epigenetic heterogeneity in the endothelial cells in ADPKD kidneys

Numerous studies have suggested that endothelial dysfunction is associated with ADPKD pathogenesis and decline of renal function through inflammation, oxidative stress, and hypoxia^38^. Endothelial cells were subclustered into 6 subtypes with distinct molecular signatures (Fig. 5a,b, Supplementary Data 17). Lymphatic vessel endothelial cells (LEC) were identified as a *PROX1*+*RELN*+ population^39^. LEC were detected primarily in ADPKD kidneys, consistent with recently reported CCL21-driven lymphangiogenesis in CKD^40^. Capillary endothelial cells (CEC), venous, and arterial endothelial cells (VEC and AEC) were detected in both ADPKD and control kidneys (Fig. 5a, b). The dominant endothelial subpopulation detected in ADPKD kidneys was *HIF1A*+*SELE*+ (EC_SELE). This subpopulation differentially expressed several cell adhesion molecules (CAM) like *SELE*, *ICAM1* and *VCAM1*(Fig. 5b,c) that have roles in recruitment of inflammatory cells^41^. This endothelial subtype was also enriched for inflammatory response gene expression (Fig. 5d). In agreement with this, RELA and AP1(JUN/FOS dimer) binding motifs were enriched on accessible chromatin in endothelial cells of ADPKD kidneys (Fig. 5e, Supplementary Data 12).

**Figure 5.**
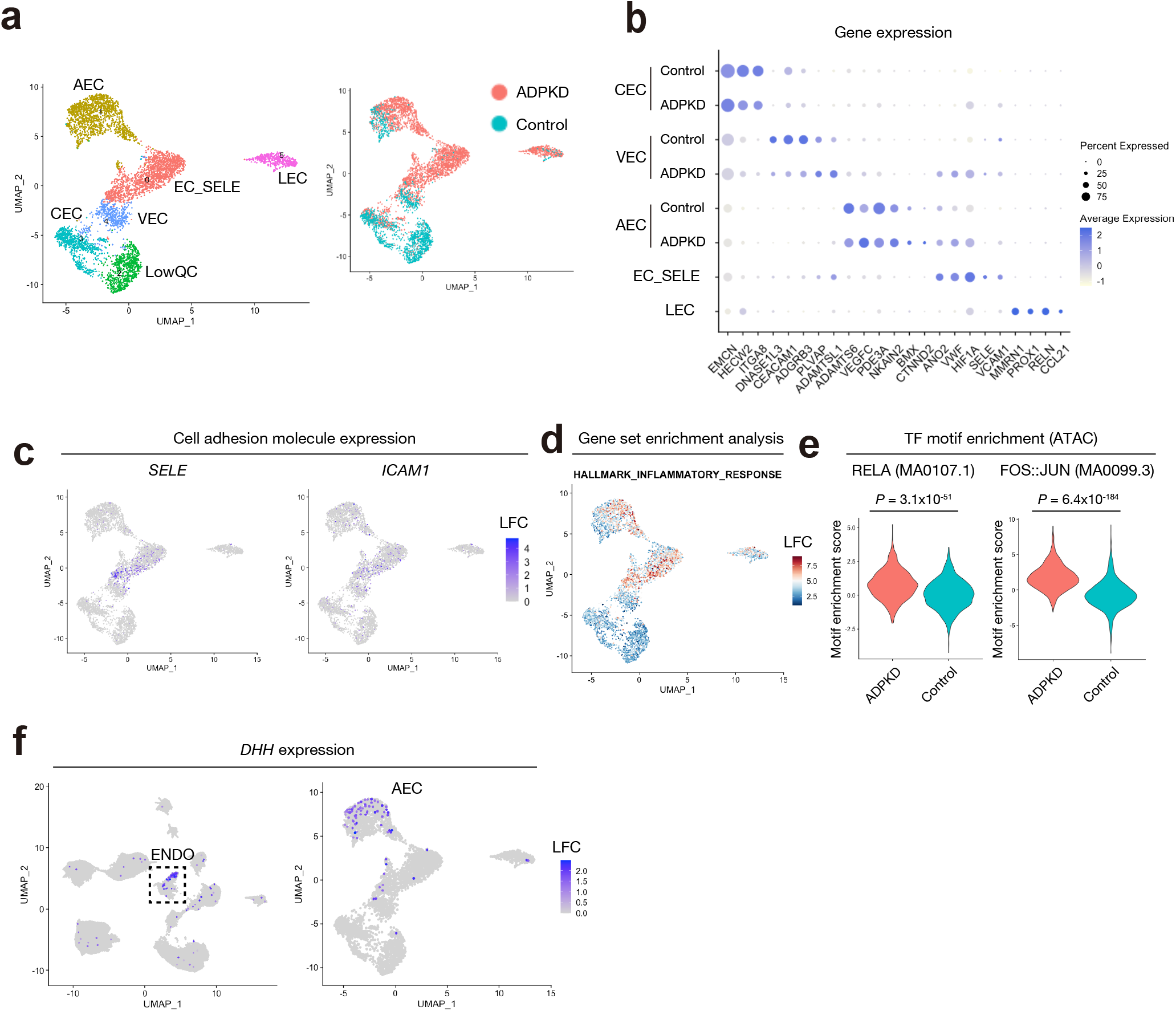
Transcriptional and epigenetic heterogeneity in the endothelial cells in ADPKD kidneys. (**a**) Subclustering of ENDO on the UMAP plot of snRNA-seq dataset with annotation by subtype (left) or disease condition (right). AEC, arterial endothelial cells; VEC, venous endothelial cells; CEC, capillary endothelial cells; EC_SELE, endothelial cells expressing *SELE*; LEC, lymphatic endothelial cells; LowQC, low-quality cells. (**b**) Dot plots showing gene expression patterns of the genes enriched in each of ENDO subpopulations. For CEC, VEC and AEC, control and ADPKD data were individually shown. The diameter of the dot corresponds to the proportion of cells expressing the indicated gene and the density of the dot corresponds to average expression relative to all ENDO cells. (**c**) Umap plot displaying *SELE* (left) or *ICAM1* (right) gene expression in the snRNA-seq. The color scale represents a normalized log-fold-change (LFC). (**d**) UMAP plot showing enrichment of gene set for inflammatory response. The color scale represents a normalized LFC. (**e**) Violin plot showing RELA (left) or AP-1 (JUN::FOS, right) motif enrichment scores for ADPKD or control kidneys. (**f**) Umap plot displaying *DHH* gene expression for the whole dataset (left) or ENDO subsets (right). The color scale represents a normalized LFC. Bonferroni adjusted p-values were used to determine significance.

We have previously shown that Gli1+ perivascular fibroblasts were adjacent to endothelial cells in mice^42^, suggesting a role of the endothelium in fibrosis. There are three Hh ligands identified in mice and in humans; sonic hedgehog (SHH), desert hedgehog (DHH), and Indian hedgehog (IHH). Interestingly, *DHH* is specifically expressed in endothelial cells, especially in AEC (Fig. 5f), suggesting that AEC may contribute to myofibroblast proliferation in ADPKD kidneys. Collectively, these findings implicate lymphangiogenesis and a proinflammatory, profibrotic molecular signature of endothelial cell subsets in ADPKD.

### Characterization of collecting duct cyst subtypes

In ADPKD kidneys, cysts are derived from both proximal and distal nephron segments. Most cyst lining cells expressed the distal nephron marker CDH1 in our samples (Fig. 3c). To characterize the cyst lining cells originated from principal cells, we performed re-clustering of the connecting tubule and principal cell (CNT_PC) cluster (Fig. 6a, Supplementary Data 18), resulting in separation to 8 clusters. Among them, two clusters exhibited enriched expression of *SLC26A7* (IC-A marker) or *LRP2* (PT marker) transcripts along with mitochondrial genes, suggesting that they were multiplets including intercalated (IC) or PT cells. Cluster1 and 2 were detected in both healthy and ADPKD kidney cells (Fig. 6a). Based on the differentially expressed genes, we annotated these as normal connecting tubules (N-CNT, cluster 1) and normal PC (N-PC, cluster2), respectively. We also detected four ADPKD-specific clusters (PKD-CNT and PKD-PC1, 2 and 3). PKD-CNT (cluster5) and PKD-PC1 (cluster0) differentially expressed *MET,* and PKD-PC2 (cluster4) differentially expressed *LCN2*. *MET* and *LCN2* were previously found to be essential for disease progression in an ADPKD mouse model^43,44^ (Fig. 6b). PKD-PC3 represented a smaller cyst subcluster with unique markers (Fig. 6b). Interestingly, ADPKD subclusters upregulated the expression of *CFTR* (Fig. 6b and Supplementary Fig. 7a,b), which encodes a chloride ion channel thought to be involved in fluid accumulation in the cyst^45^. To characterize the molecular signatures of these PKD-specific subtypes, we performed gene set enrichment analyses. This revealed general down-regulation of genes related to oxidative phosphorylation in the ADPKD clusters, and enrichment of glycolysis gene signatures in PKD-PC2 (Fig. 6c). Inflammatory gene expression was also enriched in PKD-PC1 and PKD-PC2 (Fig. 6d). These findings are consistent with the previously observed mitochondrial dysfunction, activation of glycolysis, and inflammation in ADPKD cysts^45,46^.

**Figure 6.**
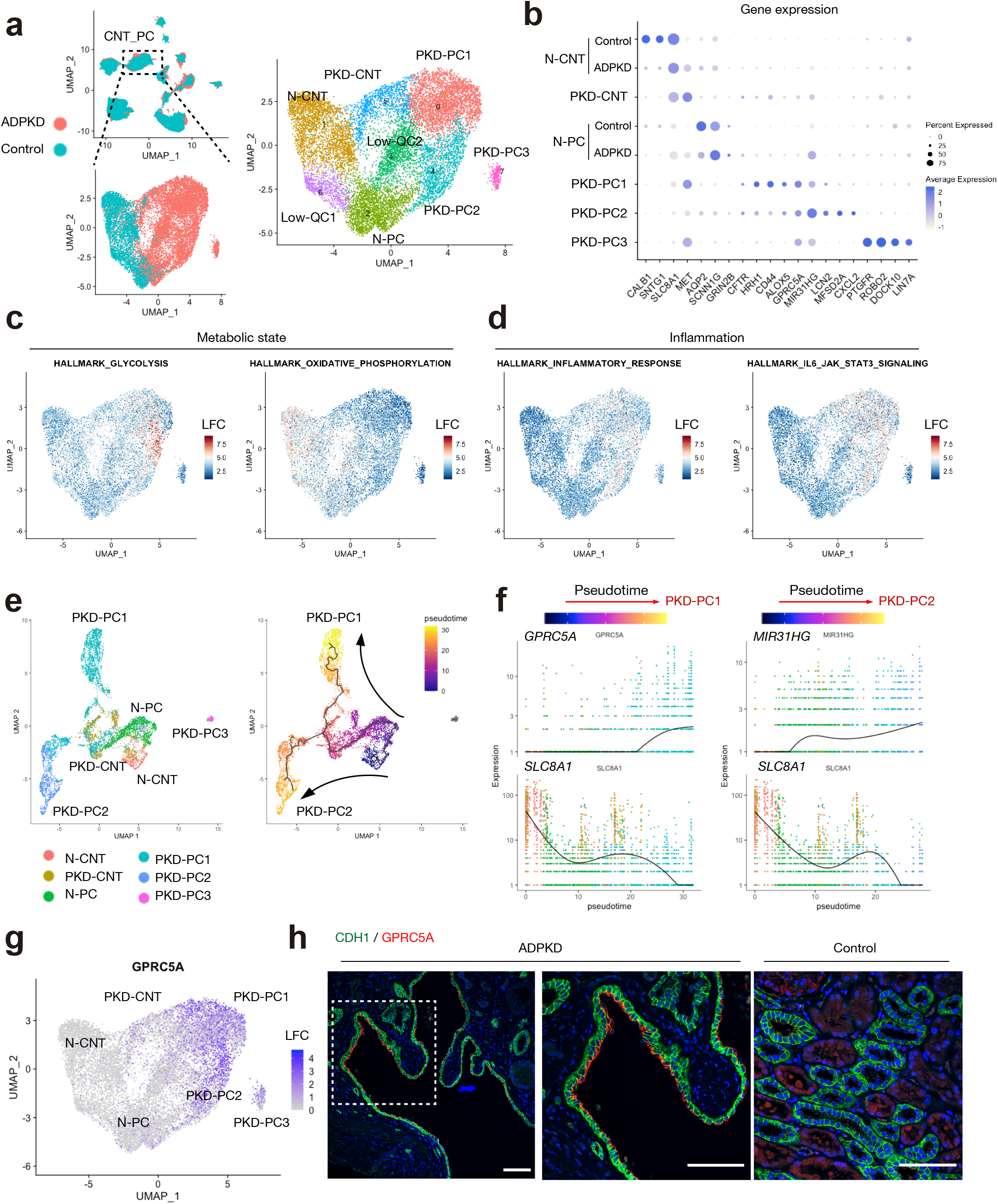
Transcriptomic characterization of cyst-lining cells originated from collecting duct. (**a**) Subclustering of CNT_PC on the UMAP plot of snRNA-seq dataset with annotation by subtype (left) or disease condition (right). N-CNT, normal CNT; N-PC, normal PC; PKD-CNT, ADPKD-specific CNT; PKD-PC, ADPKD-specific PC; LowQC, low-quality cells. (**b**) Dot plots showing gene expression patterns of the genes enriched in each of subpopulations. For N-PC and N-CNT, control and ADPKD data were individually shown. The diameter of the dot corresponds to the proportion of cells expressing the indicated gene and the density of the dot corresponds to average expression relative to all CNT_PC cells. (**c**) UMAP plot showing enrichment of gene set for metabolic states. Gene signature related to glycolysis (left) and oxidative phosphorylation (right). The color scale represents a normalized log-fold-change (LFC). (**d**) UMAP plot showing enrichment of gene set for inflammation. Gene signature related to inflammatory response (left) and IL6-induced JAK-STAT3 pathway (right). The color scale represents a normalized LFC. (**e**) Pseudotemporal trajectory in CNT_PC subpopulations of ADPKD kidneys from normal to ADPKD-specific subtypes using snRNA-seq was generated with Monocle3 (right). The subtype annotations were also shown (left). (**f**) Gene expression dynamics along the pseudotemporal trajectory toward PKD-PC1 or PKD-PC2 are shown. For PKD-PC1 branch; *GPRC5A* (upper left), *SLC8A1* (lower left), and for PKD-PC2 branch; *MIR31HG* (upper right), *SLC8A1* (lower right). (**g**) UMAP plot displaying *GPRC5A* gene expression in CNT_PC subtypes. The color scale represents a normalized LFC. (**h**) Representative immunohistochemical images of CDH1 (green) and GPRC5A (red) in the ADPKD (left and middle, n=3) or control kidneys (right, n=3). Scale bar indicates 50 μm.

Pseudotemporal trajectories were generated to show the potential differentiation pathways from normal to PKD-specific clusters (Fig. 6e). PKD-PC3 does not clearly have a lineage relationship with the other principal cells since it forms a distinct cluster so it was not included in this analysis. Interestingly, there were divergent trajectories toward PKD-PC1 and PKD-PC2. The normal CNT/PC marker gene (*SLC8A1*) was down-regulated and unique marker genes (*GPRC5A* or *MIR31HG*) were gradually up-regulated during the progression of pseudotime to PKD-PC1 or PKD-PC2 (Fig. 6e). Among the differentially expressed genes in PKD-PC1, *GPRC5A* was one of the most differentially up-regulated genes in PKD-PC1 cluster (*P adj.* < 1.0 ×10^−300^, Log fold change = 1.62, Fig. 6f,g, Supplementary Data 18). Immunofluorescence studies validated GPRC5A expression in CDH1+ cyst lining cells (Fig. 6h, left), while its expression was faint in CDH1+ tubular cells in control kidneys (Fig. 6h, right). GPRC5A expression was recently found to be up-regulated by hypoxia, and to suppress the Hippo pathway, promoting survival of the cells *in vitro*^47^. Thus, GPRC5A might confer resistance to hypoxia in cyst cells, promoting cyst growth in ADPKD.

Another interesting gene up-regulated in PKD-PC1 (*P adj.* = 2.7 ×10^−14^, Log fold change = 0.30) and PKD-PC2 (*P adj.* = 8.0 ×10^−234^, Log fold change =1.19, Supplementary Data 18) was *MIR31HG* that encodes a long non-coding RNA (Supplementary Fig. 8a). *MIR31HG* expression was found to be induced by hypoxia to promote HIF1A-dependent gene expression as a co-activator^48^, consistent with up-regulation of glycolysis-related genes in PC subpopulations of ADPKD kidneys (Fig. 6c). Furthermore, *MIR31HG* was shown to prevent cellular senescence via suppression of *CDKN2A* transcription through recruitment of polycomb group proteins^49^. Previously published studies conclude that senescence attenuates disease progression in a mouse model of ADPKD^50^. This suggests that *MIR31HG* may promote cyst growth through suppression of senescence and adaptation to hypoxia. While *CDKN1B* and *CDKN1C* expression was higher in normal subtypes (Supplementary Fig.8b), senescence-related CDK inhibitors *CDKN1A* and *CDKN2A* were more abundant in PKD-PC subtypes (Supplementary Fig. 8a,b), suggesting that cyst lining cells may be prone to cellular senescence due to cellular stress. Upregulated *MIR31HG* expression in ADPKD kidneys (Supplementary Fig.8a) could circumvent cellular senescence by inhibiting further up-regulation of *CDKN2A*.

### Long-range genomic cis-interactions govern molecular signatures of cyst lining cells

Next, we analyzed CNT_PC cluster in the snATAC-seq dataset to dissect the epigenetic mechanism driving unique molecular signatures in ADPKD cyst cells. We detected an ADPKD-specific PC subpopulation (PKD-PC, Fig. 7a, Supplementary Data 19). This subtype showed higher gene activity of *GPRC5A* or *CD44* compared to other subpopulations (Fig. 7b), suggesting that they represent the combined PKD-PC1 and PKD-PC2 subclusters previously identified in snRNA-seq data. Transcription factor motif enrichment analysis indicated activation of NF-κB transcription factors and transcriptional enhanced associate domain (TEAD) family transcription factors in PKD-PC (Fig.7c, Supplementary Data 20). TEAD family transcription factors have been shown to play important roles in tissue homeostasis and organ size control, with activity regulated by their coactivator; YAP and TAZ in the Hippo pathway^51^. Recently, ROR1 was shown to be a co-receptor of Frizzled 1, activating YAP/TAZ by WNT5A/WNT5B^52^. Furthermore, ROR1 was found to mediate WNT5A-induced NF-κB activation^53,54^. Interestingly, we observed ROR1 protein expression in GPRC5A+ cyst cells (Fig. 7d, Supplementary Fig. 9a). ROR1 expression in bulk tissue was directly correlated with cyst size in a reanalysis of published datasets^55^ (Supplementary Fig. 9b). In agreement with this, *ROR1* expression was generally up-regulated in ADPKD in our dataset (Supplementary Fig. 9c), although it not in a cell type specific fashion (Supplementary Fig. 9d). Collectively, these findings suggest a potential role of ROR1 and GPRC5A in defining a unique gene signature of cyst lining cells in ADPKD.

**Figure 7.**
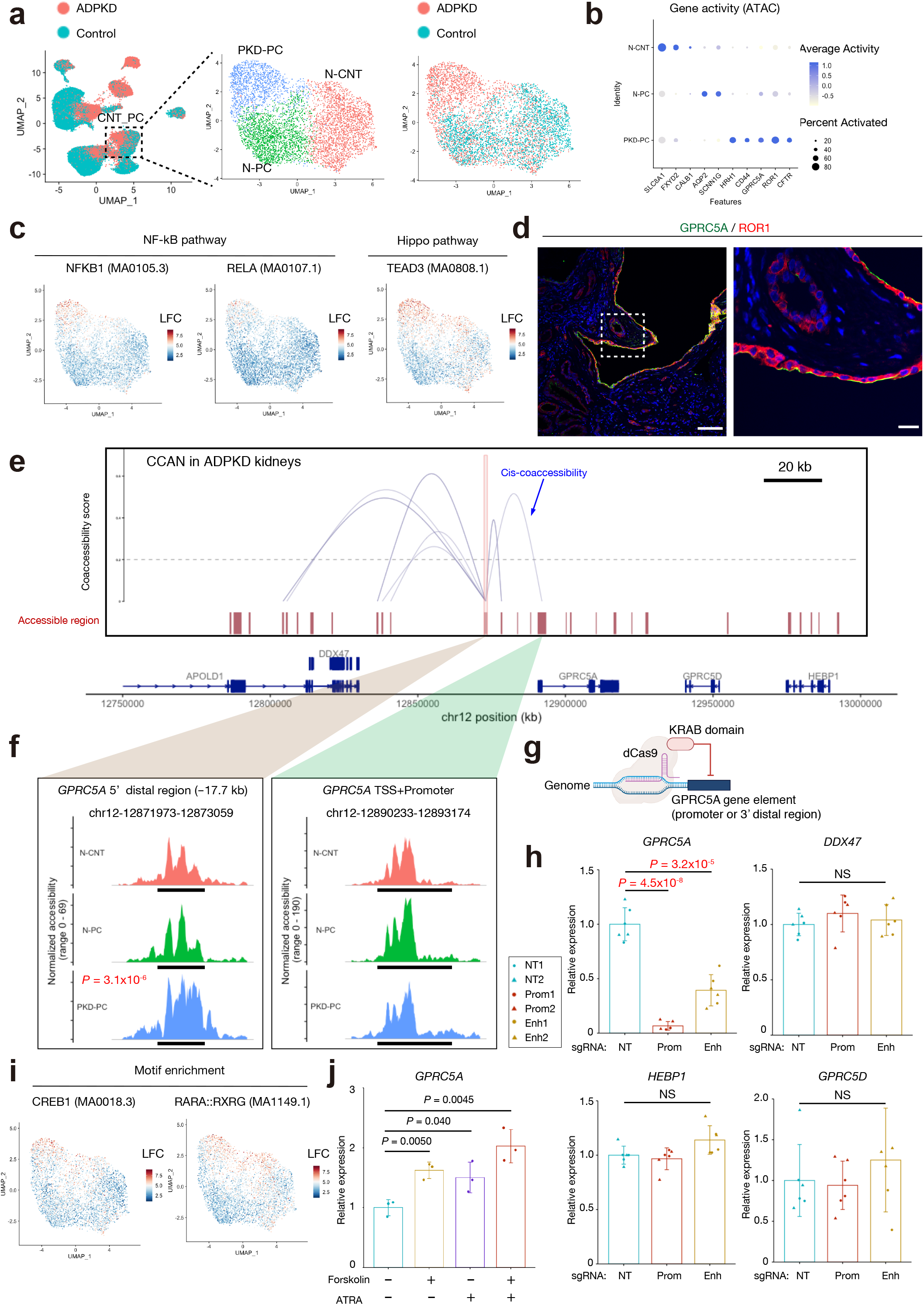
Multimodal approach revealed epigenetic alterations in ADPKD cyst cells. (**a**) Subclustering of CNT_PC on the UMAP plot of the snATAC-seq dataset with annotation by subtype (left) or disease condition (right). N-CNT, normal CNT; N-PC, normal PC; PKD-PC, ADPKD-specific PC. (**b**) Dot plots showing gene activity patterns of the genes enriched in each of CNT_PC subpopulations. The diameter of the dot corresponds to the proportion of cells with detected activity of indicated gene and the density of the dot corresponds to average gene activity relative to all CNT_PC nuclei. (**c**) UMAP plot showing enrichment of transcription factor binding motifs for NF-κB pathway; NFKB1 (left) and RELA (middle), or Hippo pathway; TEAD3 (right). The color scale represents a normalized log-fold-change (LFC). (**d**) Representative immunohistochemical images of ROR1 (red) and GPRC5A (green) in the ADPKD kidneys (n = 3). Scale bar indicates 50 μm (left) or 10 μm (right). (**e**) Cis-coaccessibility networks (CCAN, gray arcs) around the *GPRC5A* locus among accessible regions (red boxes) is shown. (**f**) Fragment coverage (frequency of Tn5 insertion) around TSS (middle right, chr12:12890233-12893174) or 5′ distal DAR (middle left, chr12:12871973-12873059) are shown (peak +/−1 Kb). Bonferroni adjusted p-values were used to determine significance for differential accessibility. (**g**) Graphic methodology showing CRISPR interference with dCas9-KRAB domain fusion protein and small guide RNAs (sgRNA) to target potential enhancer or promoter of *GPRC5A* gene. Schematic was created with BioRender. (**h**) RT and real-time PCR analysis of mRNAs for *GPRC5A* or its surrounding genes (*DDX47*, *HEBP1* and *GPRC5D*) in the primary renal proximal tubular epithelial cells (primary RPTEC) with CRISPR interference targeting the promoter (Prom) or 5′ distal potential enhancer (Enh) for *GPRC5A* gene. NT, non-targeting control. Each group consists of n = 6 data (2 sgRNAs with 3 biological replicates). Bar graphs represent the mean and error bars are the s.d. P values are based on Student’s t test. (**i**) UMAP plot showing enrichment of transcription factor binding motifs for CREB1 (left) or retinoic acid receptor (RARA::RXRG, right). The color scale represents a normalized LFC. (**j**) RT and real-time PCR analysis of mRNAs for *GPRC5A* in primary RPTEC treated with forskolin (10 μM) with or without all-trans retinoic acid (ATRA, 1 μM) for 6 h (n=3 biological replicates). Bar graphs represent the mean and error bars are the s.d. P values are based on Student’s t test.

To characterize epigenetic mechanisms driving cyst growth, a cis-coaccessibility network (CCAN) in ADPKD kidneys around *MIR31HG* or *GPRC5A* gene was predicted with Cicero^9^ (Fig. 7e, Supplementary Fig. 8c). Although *MIR31HG* is differentially expressed in ADPKD cells (Fig. 6g), the promoter region of *MIR31HG* was more accessible in normal PC cells. However, we found that the 5′ distal region to *MIR31HG* gene was differentially accessible in the ADPKD subtype, and co-accessible to the *MIR31HG* promoter (Supplementary Fig.8c). This 5′ distal region was previously shown to be an enhancer which up-regulates *MIR31HG* expression in oncogenic cellular stress^49^. Specific activation of this enhancer and *MIR31HG* upregulation in the ADPKD subtype is consistent with the notion that *MIR31HG* may promote cyst growth via suppression of cellular senescence. For the *GPRC5A* CCAN, the TSS of *GPRC5A* was not differentially accessible among subtypes. However, we identified a ~17 kb 5′ distal region that was differentially accessible in PKD-PC subpopulation was predicted to interact in cis with the *GPRC5A* TSS (Fig. 7e, Supplementary Data 19). To determine whether this 5′ distal region has an enhancer activity for *GPRC5A* gene expression, we performed CRISPR interference on the 5′ distal region in human primary kidney proximal tubular cells (primary RPTEC)^15^. While CRISPR interference on the promoter region achieved ~90% decrease of *GPRC5A* expression, targeting the 5′ distal region induced a 50~60% decrease, confirming its enhancer activity (Fig. 7f, g). Of note, that enhancer activity was specific to *GPRC5A* gene expression and not neighboring genes (Fig. 7f), consistent with its specific interaction with *GPRC5A* promoter predicted by the CCAN (Fig. 7e).

This 5′ distal enhancer has several binding motifs for cAMP responsive element binding protein 1 (CREB1) and retinoic acid receptors (RAR) as well as NF-κB (RELB) and TEAD family transcription factors (TEAD1-4), based on JASPAR 2018^56,57^ (Enrichment score > 300, Supplementary Fig. 10). Aberrant activation of cAMP signaling has been linked to disease progression in ADPKD, and CREB1 mediates cAMP-dependent gene regulation. Recent lines of evidence suggest that the retinoic acid signaling pathway mediated by RAR in the collecting duct plays a protective role in kidney injury^58^. Interestingly, retinoic acid signaling was shown to be also activated by vasopressin in collecting duct cells^59^, providing evidence of crosstalk between cAMP signaling and retinoic acid signaling pathways. The binding motifs for CREB1 and retinoic acid receptors were enriched in ADPKD cells (Fig. 7f, Supplementary Data 12), suggesting that these transcription factors were activated, potentially also inducing *GPRC5A* expression in ADPKD. In agreement with this finding, the expression of retinol dehydrogenase 10 (*RDH10*), which is a rate-limiting enzyme of retinoic acid synthesis was maintained among PC subpopulations in ADPKD kidneys (Supplementary Fig. 11). Previous literature suggests that cAMP and retinoic acid stimulation upregulates GPRC5A expression in cancer cell lines^60,61^. To validate these findings, we treated human primary RPTEC with cAMP-inducing agent forskolin with or without all trans retinoic acid, and we confirmed that cAMP and retinoic acid signaling increased *GPRC5A* expression in the kidney cell line (Fig. 7g). Although these are proximal tubule rather than principal cells, the results are consistent with the notion that cAMP drives expression of GPRC5A, potentially participating in proliferation of cyst cells.

## Discussion

We performed multimodal single cell analysis on adult human ADPKD kidneys to untangle cellular complexity and dissect the molecular foundation of disease progression at single cell resolution. Our analysis elucidates previously unrecognized cellular heterogeneity that was defined with gene expression signatures and chromatin accessibilities in ADPKD kidneys.

Previous studies in mice have described a maladaptive repair cell state in proximal tubular cells. *Vcam1*+*Ccl2*+ failed repair cell states have been identified in snRNA-seq of mouse kidneys after ischemia reperfusion injury, and their transcriptomic signatures indicated NF-κB activation^18,19^. Furthermore, we recently identified a VCAM1+ subpopulation in PT of healthy human kidneys through multimodal single cell analysis^12^. The transcriptomic signature of VCAM1+PT was shown to be similar to mouse failed repair PT population, suggesting that a small number of failed repair PT cells also exist in human healthy kidneys. In this study, we found that most of the normal PTC were replaced by a FR-PTC population in ADPKD kidneys (Fig. 3b,c). Although most of FR-PTC did not compose cyst lining cells (Fig. 3c), they were found to express proinflammatory (*CCL2*) or profibrotic (*TGFB2*) molecules (Fig. 3b), suggesting their proinflammatory and profibrotic roles in the microenvironment of the ADPKD kidney. CCL2 was previously shown to promote cystogenesis via recruitment of macrophages in model mice^62^. The role of TGFβ signaling in ADPKD progression is still controversial^63,64^, although its profibrotic effect on the kidneys has been widely accepted^31^. There have been lines of evidence indicating that cystogenesis was promoted by ischemic renal injuries in a mouse model of ADPKD^65^. FR-PTC may have a role in acceleration of cyst growth in such conditions. Interestingly, FR-PTC also up-regulated TGFβ receptor expressions, suggesting autocrine of TGFβ in FR-PTC. Autocrine loop of TGFβ signaling was previously found to lead to aberrantly high levels of TGFβ2 through CREB1 and SMAD3 binding to the *TGFB2* promoter in glioblastoma^66^. This suggests that TGFβ signaling in FR-PTC may not only induce fibrosis but also maintain FR-PTC cell state in CKD kidneys. These findings also suggested a potential therapeutic angle for cAMP-CREB1-TGFB2 axis in FR-PTC.

ADPKD is known to be associated with various degrees of interstitial fibrosis, which is the final common pathway of CKD regardless of etiology^31^. Interstitial fibrosis has been shown to be an accelerator of disease progression in ADPKD, as well as other CKD^67^. Renal fibrosis is a complicated series of processes including tissue injury, inflammation, myofibroblast proliferation, and deposition of extracellular matrix in the tissue to cause irreversible tissue remodeling^31^. Many cell types and fibroblast subtypes are involved in fibrosis, although the heterogeneity of fibroblasts in ADPKD kidneys and the roles of each fibroblast subtypes have not been elucidated. Here, we identified two fibroblast subtypes predominant in ADPKD kidneys: *ACTA2*+myofibroblasts and *FGF14*+*IL6*+fibroblasts. Interestingly, we found the latter fibroblast subtype expressed *IL6* at the highest level among all ADPKD kidney cell types (Supplementary Fig.6a,b), suggesting their role in the inflammatory microenvironment. *GLI1* expression was specifically up-regulated in these fibroblast subtypes in ADPKD. We previously showed Gli1 as a marker of myofibroblast progenitors in mouse kidney as well as other organs^42^, and Gli2 inhibition prevents proliferation, decreasing fibrosis in mouse kidneys^35^. The upregulation of *GLI1* expression observed in ADPKD fibroblast subtypes suggested activated Hh pathway, further supported by up-regulation of PTCH co-receptors *GAS1*, *CDON* or *BOC* in these subtypes (Fig.4g). Inhibition of Hh pathway may be another potential therapeutic target for ADPKD progression.

Vascular endothelial cells have been also found to be involved in ADPKD pathophysiology^68^. Vascular phenotypes, including leakage and rupture of blood vessels were described in mice with *Pkd1* or *Pkd2* deletion that brought embryonic lethality^69,70^. Although intracranial aneurysm and arterial dissection are of the most clinical interest^68^, evidence implicates endothelial dysfunction in promoting cyst growth in ADPKD^38^. We identified an inflammatory endothelial subset that expresses several cell adhesion molecules, suggesting a role in inflammatory cell recruitment. The NF-κB pathway initiated by TNFα was shown to induce such adhesion molecules in the endothelium^71^. Indeed, we found RELA transcription factor motifs were enriched on the open chromatin regions of ADPKD endothelial cells (Fig.5e), consistent with the inflammatory milieu surrounding endothelial cells. We also found that *DHH* was highly expressed in AEC (Fig. 5f), possibly involving paracrine DHH signaling from AEC to surrounding pericytes and fibroblasts for Hh pathway activation. Dhh was shown to have a protective role in ischemic injuries^72^in mice, although lines of evidence suggested aberrant activation of the Hh pathway promotes tissue remodeling and fibrosis^34^. DHH from AEC under chronic tissue hypoxia may be responsible for Hh pathway activation observed in ADPKD fibroblasts. However, further investigation is needed to address the role of DHH in cystogenesis.

Principal cells of the collecting duct have been proposed as the main origin of cyst in ADPKD^1,73,74^. We identified a GPRC5A+CDH1+ PC lineage (PKD-PC1) in cyst lining cells (Fig. 6g,h). This subpopulation also differentially expresses *MET*, which mediated HGF-dependent mTORC activation in a mouse model of ADPKD^43^. GPRC5A expression has been shown to be induced by hypoxia in a cancer cell line, subsequently activating YAP/TAZ to promote survival of the cells in hypoxic conditions^47^. YAP/TAZ-dependent gene regulation is mediated by TEAD family transcription factors^75^. In agreement with this, TEAD3 activity was increased in the PC lineage in ADPKD kidneys in snATAC-seq (Fig. 7c) Furthermore, another YAP/TAZ activator ROR1 was also expressed in GPRC5A+ cysts (Fig. 7d). ROR1 is receptor tyrosine kinase that is overexpressed in many types of cancers to activate non-canonical WNT signaling pathway. It is tempting to speculate that up-regulation of GPRC5A and ROR1 in cyst lining cells might drive proliferation in the hypoxic milieu in ADPKD kidneys. Another PC subpopulation (PKD-PC2) expresses *LCN2* that was shown to be expressed in cyst cells of a mouse model and human samples^44^. Interestingly, pseudotemporal analysis suggested both PKD-PC1 and PKD-PC2 independently arose from the normal PC/CNT lineage (Fig. 6e). PKD-PC2 expressed HIF-dependent glycolysis genes (Fig. 6c), suggesting that differences in responses to hypoxia may switch the trajectories to PKD-PC1 and PKD-PC2.

We leveraged snATAC-seq data to predict an enhancer element for GPRC5A expression in ADPKD kidneys (Fig.7), and validated that the enhancer regulates GPRC5A expression by CRISPR interference (Fig. 7h). Given that promoter accessibility was not changed between ADPKD and control cells (Fig. 7f), this differentially accessible enhancer may be responsible for up-regulation of *GPRC5A* gene expression in ADPKD PC. We observed RAR or CREB1 binding sites in that enhancer region (Supplementary Fig. 10), and these transcription factor binding motifs were enriched in accessible regions of PC lineage in ADPKD kidneys (Fig. 7i), suggesting that *GPRC5A* is regulated by retinoic acid signaling and cAMP signaling pathways. Indeed, we confirmed that *GPRC5A* expression was regulated by these signaling pathways in human primary renal tubular cells (Fig.7j). Several lines of evidence suggests retinoic acid signaling pathway in collecting duct plays a protective role against kidney injury^58^. Aberrant activation of the retinoic acid signaling pathway may promote rather cyst growth and interstitial remodeling in ADPKD kidneys. This pathway was also found to be activated by vasopressin^59^, a pathway whose inhibition slows progression in ADPKD.

*MIR31HG* was found to be overexpressed both in PKD-PC1 and PKD-PC2, although *MIR31HG* transcripts were more abundant in PKD-PC2 (Fig. 6b, Supplementary Fig. 8b). *MIR31HG* is a long non-coding RNA that regulates HIF1A-dependent transcription^48^. Highly expressed glycolysis genes in PKD-PC2 may be regulated by *MIR31HG*. Furthermore, *MIR31HG* is up-regulated in oncogene-induced cellular stress through 5′ distal enhancer activation, and in that context suppressed expression of *CDKN2A* via recruitment of polycomb group proteins^49^. Indeed, this 5′ distal enhancer was differentially accessible in ADPKD subtype in snATAC-seq (Supplementary Fig.8c). *CDKN2A* was highly expressed in PC subpopulations in ADPKD, especially in PKD-PC1 (Supplementary Fig.8b), implicating cellular stress inducing senescence on these cells. *MIR31HG* expression in PKD-PC1 or PKD-PC2 may have a role in cyst growth through suppression of cellular senescence, given the lines of evidence suggest senescence delays cyst growth^50^. Collectively, these findings shed light on heterogeneity of cyst lining cells and several potential therapeutic targets for ADPKD.

Here, we performed multimodal single cell analysis for detection of disease-specific cell states and dissection of complicated pathophysiology involving numerous cell types. Our study is limited by the fact that these human kidney samples were from end stage disease, however this is the only human source of ADPKD tissue available. Future comparison of our dataset with preclinical ADPKD models should be informative. In conclusion, our single cell multimodal analysis of human ADPKD kidney redefines cellular heterogeneity in ADPKD kidneys, and provides a single cell multiomic foundation on which to base future efforts, including spatial technologies.

## Acknowledgments

These experiments were funded by a sponsored research agreement from Chinook Therapeutics and The Baltimore Polycystic Kidney Disease Research and Clinical Core Center (NIDDK, P30DK090868). Additional support was from the Japan Society for the Promotion of Science (JSPS) Postdoctoral Fellowships for Research Abroad and The Osamu Hayaishi Memorial Scholarship for Study Abroad (Y.M)

## Author contributions

Y.M. and B.D.H. conceived, coordinated, and designed the study. Y.M. performed experiments with contributions from E.E.D., Y.Y. and K.O. Y.M., H.W., and E.O. performed bioinformatic analysis. Y.M., E.E.D., Y.Y., H.W., A.J.K., E.O., M.G., J.K., P.A.W., T.J.W., K.O., J.H.M. and B.D.H. analyzed data. ADPKD samples were collected by E.E.D., S.L.S., O.M.W. and T.J.W. Y.M. and B.D.H. designed experiments and wrote the manuscript. All authors read and approved the final manuscript.

## Methods

### Tissue procurement

ADPKD kidney cortical cup samples were obtained from patients undergoing simultaneous native nephrectomy and living donor kidney transplantation at the University of Maryland Medical Center (Baltimore, MD). All participants provided written informed consent for participation and tissue donation. The human subjects protocol was approved by the Institutional Review Board of the University of Maryland, Baltimore. Non-tumor kidney cortex samples for controls were obtained from patients undergoing partial or radical nephrectomy for renal mass at Brigham and Women’s Hospital (Boston, MA) under an established Institutional Review Board protocol approved by the Mass General Brigham Human Research Committee. The multimodal single cell dataset generated from control kidneys (10X Genomics Chromium Single Cell 5’ v2 chemistry and 10X Genomics Chromium Single Cell ATAC v1) were already published^12^. Control kidneys were newly processed to obtain snRNA-seq libraries with 10X Genomics Chromium Single Cell 3’ v3 chemistry for this manuscript. All participants provided written informed consent in accordance with the Declaration of Helsinki. Samples were frozen or retained in paraffin blocks for future studies.

### Nuclear dissociation for library preparation

For snATAC-seq, nuclei were isolated with Nuclei EZ Lysis buffer (NUC-101; Sigma-Aldrich) supplemented with protease inhibitor (5892791001; Roche). Samples were cut into < 2 mm pieces, homogenized using a Dounce homogenizer (885302–0002; Kimble Chase) in 2 ml of ice-cold Nuclei EZ Lysis buffer, and incubated on ice for 5 min with an additional 2 ml of lysis buffer. The homogenate was filtered through a 40-μm cell strainer (43–50040–51; pluriSelect) and centrifuged at 500 g for 5 min at 4°C. The pellet was resuspended, washed with 4 ml of buffer, and incubated on ice for 5 min. Following centrifugation, the pellet was resuspended in Nuclei Buffer (10× Genomics, PN-2000153), filtered through a 5-μm cell strainer (43-50005-03, pluriSelect), and counted. For snRNA-seq preparation, the RNase inhibitors (Promega, N2615 and Life Technologies, AM2696) were added to the lysis buffer, and the pellet was ultimately resuspended in nuclei suspension buffer (1x PBS, 1% bovine serum albumin, 0.1% RNase inhibitor). Subsequently, 10X Chromium libraries were prepared according to manufacturer protocol.

### Single nucleus RNA sequencing and bioinformatics workflow

Eight ADPKD and five control snRNA-seq libraries were obtained using 10X Genomics Chromium Single Cell 3’ v3 chemistry following nuclear dissociation. A target of 10,000 nuclei were loaded onto each lane. The cDNA for snRNA libraries was amplified for 15 cycles. Libraries were sequenced on an Illumina Novaseq instrument and counted with cellranger v6.0.0 with --include-introns argument using GRCh38. The read configuration for the libraries was 2×150bp paired-end. A mean of 408,304,417 reads (s.d. = 342,469,382, control) or 358,474,996 reads (s.d. = 59,365,096, ADPKD) were sequenced for each snRNA library corresponding to a mean of 33,629 reads per cell (s.d.= 23,330, control) or 44,799 reads per cell (s.d.= 32,727, ADPKD, Supplementary table 4). The mean sequencing saturation was 47.6 +/− 13.4% (control) or 53.0 +/− 15.4% (ADPKD, Supplementary table 5). The mean fraction of reads with a valid barcode (fraction of reads in cells) was 53.9 +/− 6.8% (control) or 36.9 +/− 5.9% (ADPKD, Supplementary table 5).

Subsequently, ambient RNA contamination was corrected for each dataset by SoupX v1.5.0^76^ with automatically calculated contamination fraction. Each of datasets was then preprocessed with Seurat v4.0.0^16^ to remove low-quality nuclei (nuclei with top 5% and bottom 1% in the distribution of feature count or RNA count, or those with %Mitochondrial genes > 0.25). Heterotypic doublets were identified with DoubletFinder v2.0.3^33^ assuming 8% of barcodes represent heterotypic doublets), and resultant estimated doublets were to be removed after merging datasets. The datasets from ADPKD or control kidneys were integrated in Seurat using the IntegrateData function with anchors identified by FindIntegrationAnchors function (Supplementary Fig. 1a, 2a). Subsequently, the doublets and low QC clusters were removed for these datasets (Supplementary Fig. 1b, 2b). The ADPKD and control datasets were integrated with batch effect correction with Harmony v1.0^17^ using “RunHarmony” function on assay “RNA” in Seurat (Fig. 1). Then, there was a mean of 8127+/− 1692 nuclei in control or 7759 +/− 3377 nuclei in ADPKD per snRNA-seq library. The number of unique molecular identifiers (UMI) per nucleus was a mean of 3536 +/− 1914 in control or 2346 +/− 1274 in ADPKD. The number of detected genes per nucleus was a mean of 2222 +/− 803 genes in control or 1743 +/− 573 genes in ADPKD. %Mitochondrial genes in a nucleus was 0.027 +/− 0.050% in control or 0.0077 +/− 0.029% in ADPKD (Supplementary Fig. 3). Clustering was performed by constructing a KNN graph and applying the Louvain algorithm. Dimensional reduction was performed with UMAP and individual clusters were annotated based on expression of lineage-specific markers (Fig. 1). The final snRNA-seq library contained 62,073 nuclei from ADPKD and 40,637 nuclei from control kidneys, and represented all major cell types within the kidney cortex (Supplementary Table 2,3 and Fig. 1). Differential expressed genes among cell types were assessed with the Seurat FindMarkers function for transcripts detected in at least 20% of cells using a log-fold-change threshold of 0.25. Differential expressed genes between ADPKD and control cells in each cell type were assessed using a log-fold-change threshold of 0.25 (Supplementary Data1-3). Bonferroni-adjusted p-values were used to determine significance at an FDR◻<◻0.05.

### Single nucleus ATAC sequencing and bioinformatics workflow

Eight ADPKD kidney snATAC-seq libraries were obtained using 10X Genomics Chromium Single Cell ATAC v1 chemistry following nuclear dissociation. Five control snATAC-seq libraries (Control 1-5) were prepared and published in a prior study (GSE151302 [https://www.ncbi.nlm.nih.gov/geo/query/acc.cgi?acc=GSE151302])^12^. Libraries were sequenced on an Illumina Novaseq instrument and counted with cellranger-atac v1.2 (10X Genomics) using GRCh38. The read configuration was 2×150bp paired-end. Sample index PCR was performed at 13 cycles. A mean of 334,652,440 reads were sequenced for each snATAC library (s.d.= 95,862,297) corresponding to a median of 21,671 fragments per cell (s.d.= 11,946, Supplementary Table 5). The mean sequencing saturation for snATAC libraries was 31.6 +/− 9.7% and the mean fraction of reads with a valid barcode was 95.2 +/− 3.9% (Supplementary Table 4). The libraries from control and ADPKD kidneys were aggregated with cellranger-atac v1.2.0. Subsequently, the aggregated dataset was processed with Seurat v4.0.0 and its companion package Signac v1.1.1^11^. Low-quality cells were removed from the aggregated snATAC-seq library (subset the high-quality nuclei with peak region fragments > 1000, peak region fragments < 12000, %reads in peaks > 15, blacklist ratio < 0.005, nucleosome signal < 3 & TSS enrichment > 2). Latent semantic indexing was performed with term-frequency inverse-document-frequency (TFIDF) followed by singular value decomposition (SVD). A KNN graph was constructed to cluster cells with the Louvain algorithm. Batch effect was corrected with Harmony^17^ using the “RunHarmony” function in Seurat. A gene activity matrix was constructed by counting ATAC peaks within the gene body and 2kb upstream of the transcriptional start site using protein-coding genes annotated in the Ensembl database. The gene activity matrix was log-normalized.

For label transfer, the above snATAC-seq Seurat object was divided to control and ADPKD kidney dataset, and label transfer was performed on each of the control and ADPKD kidney dataset, using filtered control and ADPKD snRNA-seq dataset (Supplementary Fig. 1b, 2b), respectively. “FindTransferAnchors” and “TransferData” functions were used for label transfer, according to instructions (https://satijalab.org/signac/)^11^. After label transfer, the control and ADPKD snATAC-seq datasets were filtered using an 80% confidence threshold for low-resolution cell type assignment to remove heterotypic doublets (Supplementary Fig. 4). The filtered control and ADPKD snATAC-seq objects were merged and reprocessed with TFIDF and SVD. Subsequently, the dataset was processed for batch effect correction with Harmony^17^, clustering and cell type annotation based on lineage-specific gene activity (Fig. 2b–d). The final snATAC-seq library contained a total of 128,008 peak regions among 50,986 nuclei (33,621 nuclei for control and 17,365 nuclei) and represented all major cell types within the kidney cortex (Supplementary Table 2). The number of fragments in peaks per nucleus was a mean of 5560 +/− 2325 in control or 4186 +/− 2196 in ADPKD, %Fragments per nucleus in reads was a mean of 57.0 +/− 10.8% in control or 38.6 +/− 11.8% in ADPKD. Fraction of reads in peaks, number of reads in peaks per cell and ratio of reads in genomic blacklist regions per cell for each patient were shown in Supplementary Fig. 3. Differential chromatin accessibility among cell types was assessed with the Seurat FindMarkers function for peaks detected in at least 20% of cells with a likelihood ratio test and a log-fold-change threshold of 0.25. Differential chromatin accessibility between ADPKD and control cells in each cell type was assessed using a log-fold-change threshold of 0.25 (Supplementary Data4-6). Differential gene activities among cell types or between ADPKD and control cells in each cell type were assessed with the Seurat FindMarkers function with a log-fold-change threshold of 0.25 (Supplementary Data7-9). Bonferroni-adjusted p-values were used to determine significance at an FDR◻<◻0.05.

### Subclustering of each cell type

For subclustering of snRNA-seq data, the target cell type was extracted based on the annotations on the integrated dataset (Fig. 1b). Subsequently, the target cell type was further filtered based on the annotations on each of control and ADPKD datasets (Supplementary Fig. 1d, Supplementary Fig. 2d) to extract the target cell type with high confidence. The batch effects in the extracted target cell type was corrected with Harmony v1.0^17^. Subsequently, clustering was performed by constructing a KNN graph and applying the Louvain algorithm. Dimensional reduction was performed with UMAP. For snATAC-seq data, the target cell type was extracted based on the annotations on the integrated dataset (Fig. 2b). PT included all subtypes of PT (PCT, PST and FR-PTC, Fig.3g, h). Subsequently, clustering was performed by constructing a KNN graph and applying the Louvain algorithm. Dimensional reduction was performed with UMAP. FindMarkers function was used to assess differentially expressed genes, differentially accessible regions or differentially enriched transcription factor binding motifs with a log-fold-change threshold of 0.25 (Supplementary Data13-20). Bonferroni-adjusted p-values were used to determine significance at an FDR◻<◻0.05.

### Estimation of transcription factor activity from snATAC-seq data

Transcription factor activity was estimated using the integrated snATAC-seq dataset and chromVAR v1.10.0^10^. The positional weight matrix was obtained from the JASPAR2018 database^56^. Cell-type-specific chromVAR activities were calculated using the RunChromVAR wrapper in Signac v1.1.1 and differential activity was computed with the FindMarkers function with mean.fxn=rowMeans and fc.name=“avg_diff”. (Log-fold-change > 0.25 for comparison among cell types and Log-fold-change > 0.1 for comparison between ADPKD and control in each cell type, Supplementary Data10-12).

### Generation of cis-coaccessibility networks with Cicero

Cis-coaccessibility networks were predicted using Cicero v1.3.4.11 according to instructions provided on GitHub (https://cole-trapnell-lab.github.io/cicero-release/docs_m3/)^9^. Briefly, the ADPKD data was extracted from integrated snATAC-seq dataset and converted to cell dataset (CDS) objects using the make_atac_cds function. The CDS object was processed using the detect_genes() and estimate_size_factors() functions with default parameters prior to dimensional reduction and conversion to a Cicero CDS object. ADPKD-specific Cicero connections were obtained using the run_cicero function with default parameters.

### Construction of pseudotemporal trajectories on snRNA-seq data

Pseudotemporal trajectories were constructed on Monocle 3 according to instructions provided on GitHub (https://cole-trapnell-lab.github.io/monocle3)^24–26^. Briefly, raw count matrix subset (PT [Fig.3] or PC without LowQC clusters [Fig.6] in ADPKD kidneys) was used to generate cell data set object (CDS) with “new_cell_data_set” function. Next, the CDS was preprocessed (num_dim = 10 [Fig. 3] or 20 [Fig. 6]), aligned to remove batch effect among patients^77^ and reduced onto a lower dimensional space with the “reduce_dimension” function (preprocess_method = “Aligned“). The cells were then clustered (“cluster_cells“) and ordered with the “learn_graph” function. We used the “order_cell” function and indicated the most distant cell from FR-PTC (Fig.3, PT) or PKD-PC1/2 (Fig. 6, PC) as “start point” of the trajectories.

### Single cell gene enrichment analysis on snRNA-seq data

Single cell gene set enrichment analysis was performed with the VISION v2.1.0 R package according to instructions provided on GitHub (https://github.com/YosefLab/VISION)^27^, using Hallmark gene sets obtained from the Molecular Signatures Database v7.4 distributed at the GSEA Web site. The resultant enrichment scores were incorporated into metadata of the Seurat object, and visualized on UMAP plot with “FeaturePlot” function.

### Pathway analysis on snRNA-seq data with PROGENy

PROGENy (v. 1.15.3, https://saezlab.github.io/progeny/) was applied to the snRNA-seq Seurat object with “progeny” function^29^. Each pathway was scaled to have a mean activity of 0 and a standard deviation of 1. The PROGENy pathway activity scores were computed on the scRNA-seq data, and then the different cell populations were characterized based on these scores. The different pathway activities for the different cell populations were then plotted as heatmaps.

### Correlation between ROR1 expression and cyst size

The human microarray dataset GSE7869 was retrieved from the Gene Expression Omnibus database (GEO)^55^. The dataset contained n=3 non-ADPKD control kidneys, n=5 of minimally cystic tissue, n=5 of small-sized renal cysts, n=5 of medium-sized renal cysts, and n=3 of large-sized renal cysts. The renal cyst samples were obtained from five *PKD1*-mutant polycystic kidneys. To visualize the correlation of ROR1 in renal cysts, the expression level of ROR1 in units of normalized signal intensity was plotted against the grouped cyst size.

### Identification of transcription factor binding motifs in *GPRC5A* enhancer

Identification of transcription factor binding motifs in a *GPRC5A* enhancer was performed on UCSC genome browser^57^ with TFBS predictions in Homo sapiens (hg38) in the JASPAR CORE vertebrates collection^56^. Minimum score was set to 300 which corresponds to P value of 0.001.

### Immunofluorescence studies

Deparaffination of tissue samples was performed by immersing glass slides into coplin jars with xylene and ethanol (5◻min in 100% xylene, 5◻min in 100% xylene, 5 min in 100% ethanol, 5◻min in 95% ethanol, 5◻min in 70% ethanol, 5◻min in distilled water and 5◻min in distilled water). Following the last wash, slides were placed in antigen retrieval solution (Vector H-330). Samples were incubated in a pressure cooker (Prestige Medical Classic 2100 series). Following this incubation, samples were allowed to cool to room temperature, and washed with distilled water. Samples were treated with 2–3 drops of Image-iT FX Signal Enhancer (Molecular Probes; 136933) for 15◻min with rotation at room temperature, and then blocked in Blocking Media [1% BSA (Roche; 03 116 956001), 0.1% Triton X-100 (Sigma; T8787), 0.1% sodium azide (Sigma; S28032) in PBS] for another 15◻min with rotation at room temperature. Primary antibody was added in Blocking Media [rabbit Anti-GPRC5A (Sigma; SAB4503536; 1:100), goat Anti-ROR1 (Abcam; Ab111174; 1:125), mouse Anti-E-Cadherin (BD Transduction; 610182; 1:200, lotus tetragonolobus lectin (LTL) (Vector Labs; B-1325; 1:100), Anti-VCAM1 (abcam; ab134047; 1:200)] and incubated overnight in a humidifier chamber at 4°C. The next day, slides were quickly washed three times in PBS. Secondary antibodies [Donkey Anti-Goat (Invitrogen; A11057; 1:200), Donkey Anti-Mouse (Invitrogen; A21202; 1:200), Donkey Anti-Rabbit (Invitrogen; A10042 or Jackson ImmunoResearch; 711-545-152; 1:200), conjugated Streptavidin (Invitrogen; S21374; 1:200)] were added in Blocking Media and incubated at room temperature, in the dark for 1 h. Slides were then again quickly washed in PBS, incubated with DAPI (Invitrogen; D1306; 1:1000) in PBS for 5 min, and washed finally with PBS two more times for 5 min each. Following washes, coverslips were mounted onto glass slides with Prolong Gold Antifade Mountant (Invitrogen; P3690), and sealed 16 h later with nail polish. Imaging was performed on a Nikon Eclipse Ti Confocal at 10X and 20X objective and processed using Nikon Elements-AR and FIJI (Version 2.0.0-rc-68/1.52k).

### Cell culture

HEK293T cells (ATCC; CRL-3216) were maintained in a humidified 5% CO2 atmosphere at 37°C in Dulbecco’s modified Eagle’s medium (DMEM, Gibco; 11965092) supplemented with 10% fetal bovine serum (Gibco; 10437028) and antibiotics. Human primary proximal tubular cells (human RPTEC, Lonza; CC-2553) were cultured with renal epithelial cell growth medium kit (Lonza; CC-3190) in a humidified 5% CO2 atmosphere at 37°C. At 50-60% confluency, human RPTECs were treated with or without 10 μM of forskolin (Selleck Chemicals; S2449) and/or 1 μM of retinoic acid (Millipore Sigma; R2625) for 6 h. Experiments were performed on early passages.

### CRISPR interference

To generate lentiviral vectors, we designed small guide RNA (sgRNA) to target *GPRC5A* promoter or the 5′ distal region coaccessible to that promoter with CHOPCHOP (https://chopchop.cbu.uib.no/). These sgRNAs and two non-targeting control sgRNAs were placed downstream of the U6 promoter on the dCas9-KRAB repression plasmid (pLV hU6-sgRNA hUbC-dCas9-KRAB-T2a-Puro, Addgene; 71236, a gift from Charles Gersbach)^15^ with golden gate assembly. The oligonucleotides incorporated into dCas9-KRAB repression plasmids are listed on the Supplementary Table 7. Cloning with golden gate assembly was performed with Esp3I restriction enzyme (NEB, R0734L) and T4 DNA ligase (NEB, M0202L) on a thermal cycler repeating 37 °C for 5min and 16°C for 5min for 60 cycles, followed by transformation to NEB 5-alpha Competent E. coli (NEB, C2987H) as manufacturer’s instruction. The cloned lentiviral vectors were purified with mini high-speed plasmid kit (IBI Scientific; IB47102), and sgRNA incorporations were confirmed with Sanger sequencing by GENEWIZ. psPAX2 (Addgene; 12260) and pMD2.G (Addgene; 12259) were cloned and purified with HiPure Plasmid Midiprep Kit (Invitrogen; K210005).

For generation of lentiviral supernatants, HEK293T cells were seeded at 6.0×10^5^ cells per well on 6-well tissue culture plates 16 h before transfection. Then, cells were transfected with 1.5 μg of psPAX2 (Addgene; 12260, a gift from Didier Trono), 0.15 μg of pMD2.G (Addgene; 12259, a gift from Didier Trono) and 1.5 μg of dCas9-KRAB repression plasmid per well by Lipofectamine 3000 transfection reagent (Invitrogen; L3000015) as the manufacturer’s instructions. Culture media were changed to DMEM supplemented with 30% FBS 24 h after transfection. Lentivirus-containing supernatants were collected 24 h later, and they were filtered with 0.45 μm PVDF filters (CELLTREAT; 229745). The resultant supernatants were immediately used for lentiviral transduction. Human RPTEC were seeded at 5.0×10^4^ cells per well on 6-well tissue culture plates 16 h before transfection. The media on human RPTEC was then changed to the fresh lentiviral supernatants supplemented with polybrene (0.5 μg/ml, Santa Cruz Biotechnology; sc-134220) and cultured for 24 h. Subsequently, RPTEC cells were cultured in DMEM with 10% FBS and puromycin (3 μg/ml, invivogen; ant-pr-1) for 72 h.

### Quantitative PCR

RNA from human RPTECs was extracted using the Direct-zol MicroPrep Plus Kit (Zymo) following the manufacturer’s instructions. The extracted RNA (1-2 μg) was reverse transcribed using the High-Capacity cDNA Reverse Transcription Kit (Life Technologies). Quantitative PCR was carried out in the BioRad CFX96 Real-Time System using iTaq Universal SYBR Green Supermix (Bio-Rad). Expression levels were normalized to *GAPDH*, and the data were analyzed using the 2-ΔΔCt method. The following primers were used: *GAPDH*: Fw 5′-GACAGTCAGCCGCATCTTCT −3′; Rv 5′-GCGCCCAATACGACCAAATC −3′; *GPRC5A*: Fw 5′-ATGGCTACAACAGTCCCTGAT −3′; Rv 5′-CCACCGTTTCTAGGACGATGC −3′; *DDX47*: Fw 5′-GCACCCGAGGAACACGATT −3′; Rv 5′-TCCATCCCAACTGGTCACAAG −3′; *HEBP1*: Fw 5′-TTGGCAGGTCCTAAGCAAAGG −3′; Rv 5′-CTTCCCGTAGAGCCTCATCC −3′; *GPRC5D*: Fw 5′-CTGCATCGAGTCCACTGGAG −3′; Rv 5′-AAGAGTAGCAGAATTGTGACCAC −3′.

### Statistical analysis

No statistical methods were used to predetermine sample size. Experiments were not randomized and investigators were not blinded to allocation during library preparation, experiments or analysis. Quantitative PCR data (Fig. 7) are presented as mean±s.d. and were compared between groups with a two-tailed one sample Student’s *t*-test. A *P* value of <0.05 was considered statistically significant.

## Data availability

All relevant data are available from the corresponding authors on reasonable request. Sequencing data is deposited in GEO under accession number GSE185948. Previously published snATAC-seq data for five control kidneys are available in GEO (GSE151302). Public data repositories used for our analyses include Ensembl http://useast.ensembl.org., Genome UCSC browser http://genome.ucsc.edu., and JASPAR http://jaspar.genereg.net.

## Code availability

No customized code was used for data analyses in this study. Analyses were following publicly available instructions from Seurat (http://satijalab.org/seurat/), Signac (https://satijalab.org/signac/) and Monocle 3 (http://cole-trapnell-lab.github.io/monocle-release/docs/).

