## Supplementary Material for "Defining cellular complexity in human autosomal dominant polycystic kidney disease by multimodal single cell analysis"

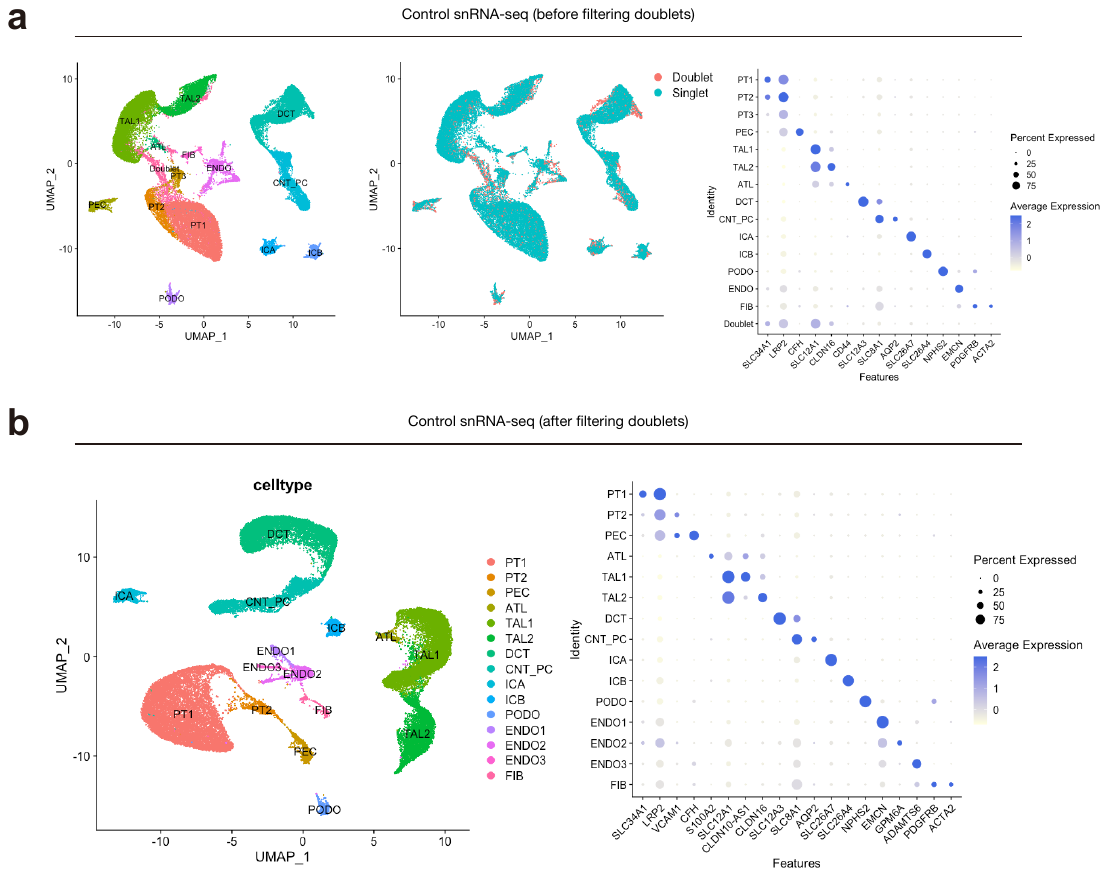

**Supplementary Figure 1. Single cell transcriptional profiling on the control kidneys**

(**a**) UMAP plots of control snRNA-seq dataset before filtering out doublets. Annotation by cell type (left) or doublet prediction by DoubletFinder (middle). Dot plot showing gene expression patterns of cluster-enriched markers (right). (**b**) UMAP plots of control snRNA-seq dataset after filtering out doublets. Annotation by cell type (left). Dot plot showing gene expression patterns of cluster-enriched markers (right). The diameter of the dot corresponds to the proportion of cells expressing the indicated gene and the density of the dot corresponds to average expression relative to all cell types.

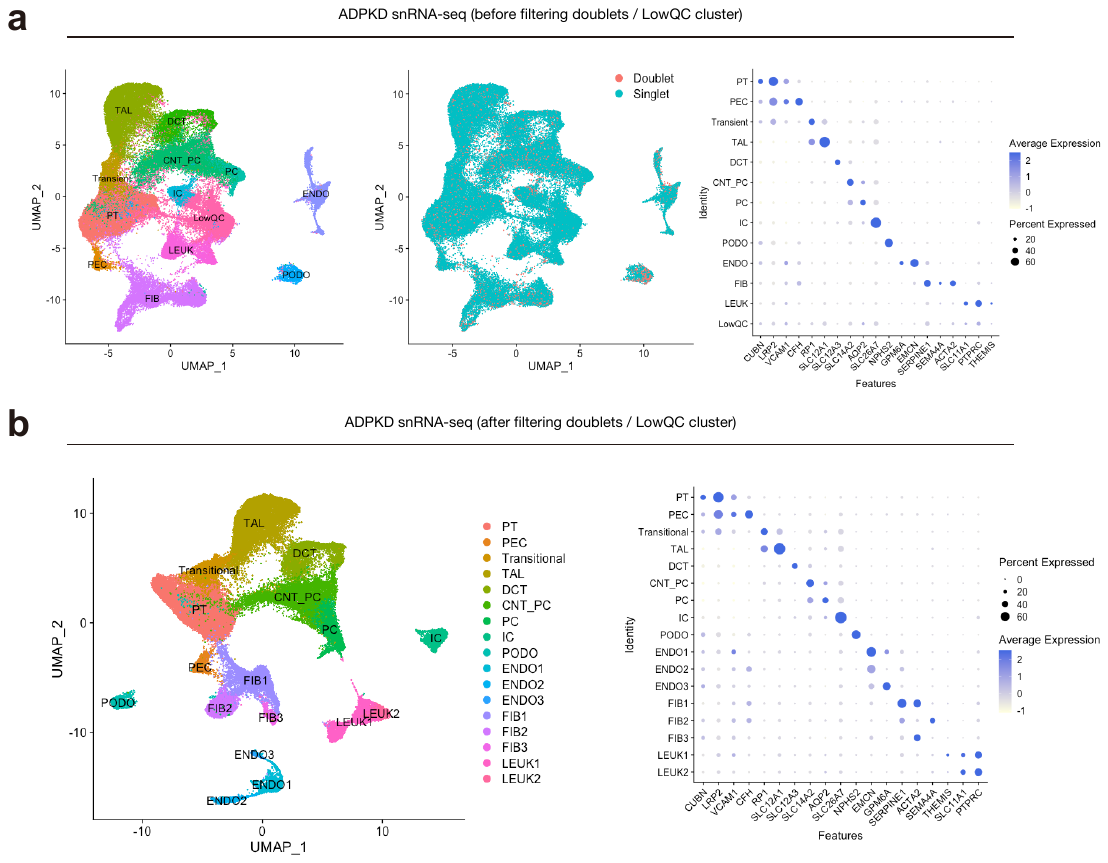

**Supplementary Figure 2. Single cell transcriptional profiling on the ADPKD kidneys**

(**a**) UMAP plots of ADPKD snRNA-seq dataset before filtering out doublets and low quality (LowQC) cluster. Annotation by cell type (left) or doublet prediction by DoubletFinder (middle). Dot plot showing gene expression patterns of cluster-enriched markers (right). (**b**) UMAP plots of ADPKD snRNA-seq dataset after filtering out doublets and LowQC cluster. Annotation by cell type (left). Dot plot showing gene expression patterns of cluster-enriched markers (right). The diameter of the dot corresponds to the proportion of cells expressing the indicated gene and the density of the dot corresponds to average expression relative to all cell types.

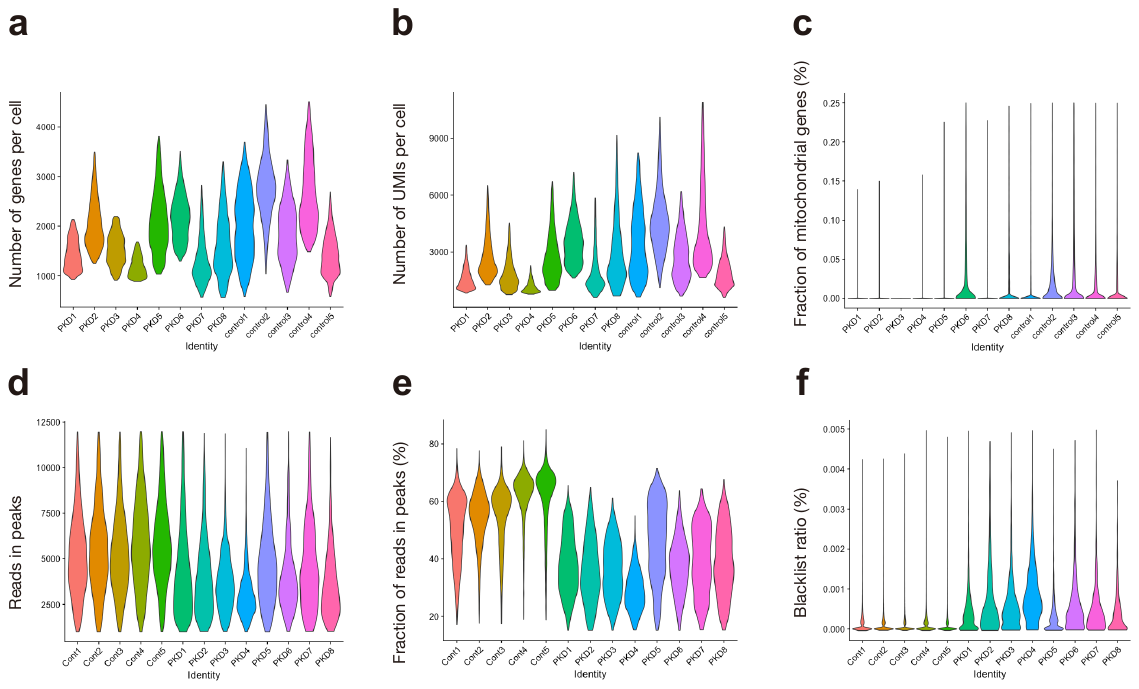

**Supplementary Figure 3. QC metrics for snRNA-seq or snATAC-seq dataset**: (**a**) Number of genes per cell, (**b**) number of UMIs per cell and (**c**) fraction of mitochondrial genes per cell in snRNA-seq data were shown. (**d**) Fraction of reads in peaks, (**e**) number of reads in peaks per cell and (**f**) ratio of reads in genomic blacklist region per cell in snATAC-seq data were shown.

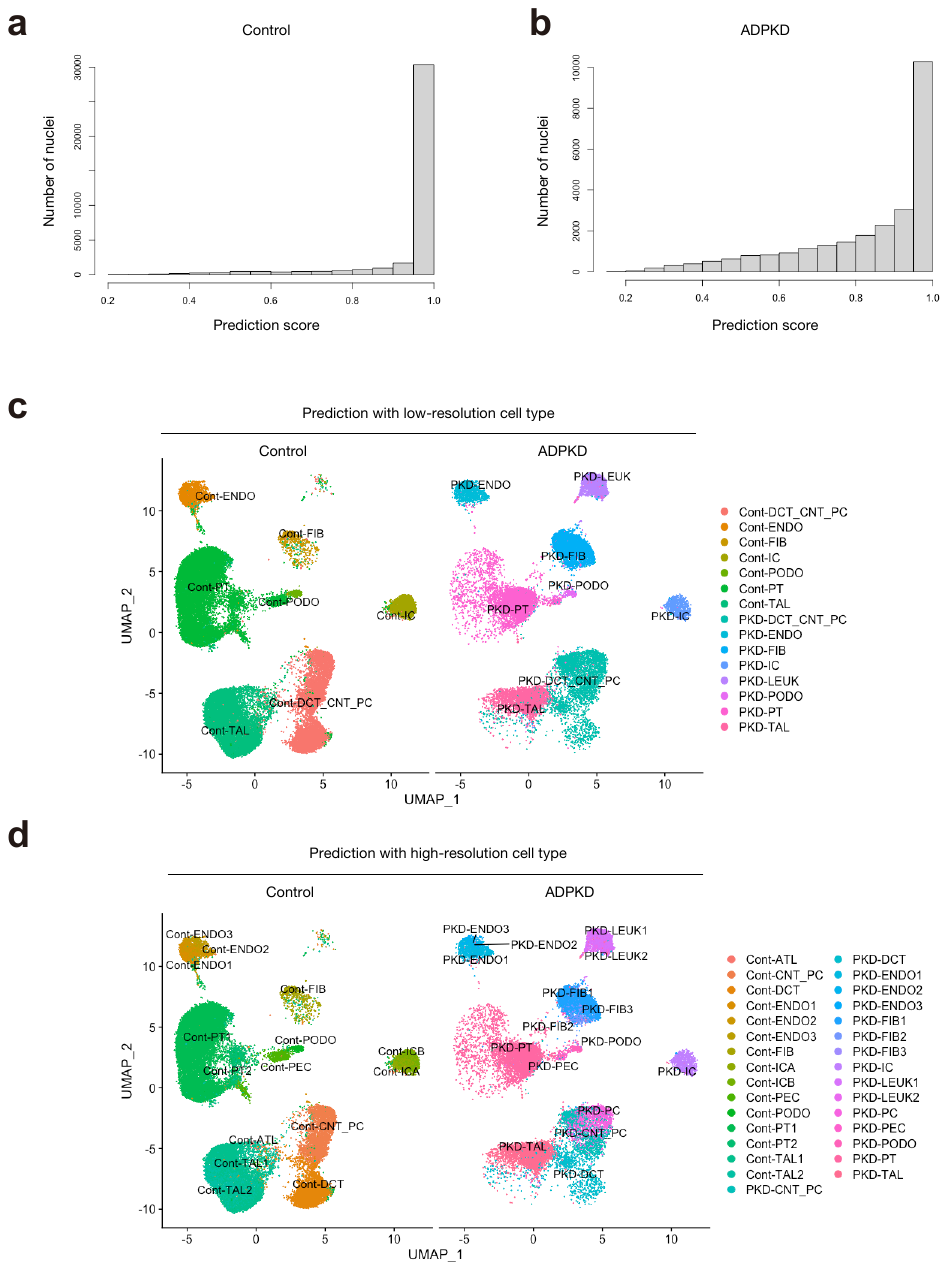

**Supplementary Figure 4. Most of nuclei in ADPKD and control kidneys were predicted with high confidence**

(**a,b**) Distribution of maximum prediction scores of nuclei calculated by the label transfer algorithm in Signac package for control (**a**) or ADPKD dataset (**b**). (**c,d**) UMAP plot of snATAC-seq dataset with predicted cell types through label transfer from snRNA-seq data with low-resolution cell types (**c**) or high-resolution cell types (**d**).

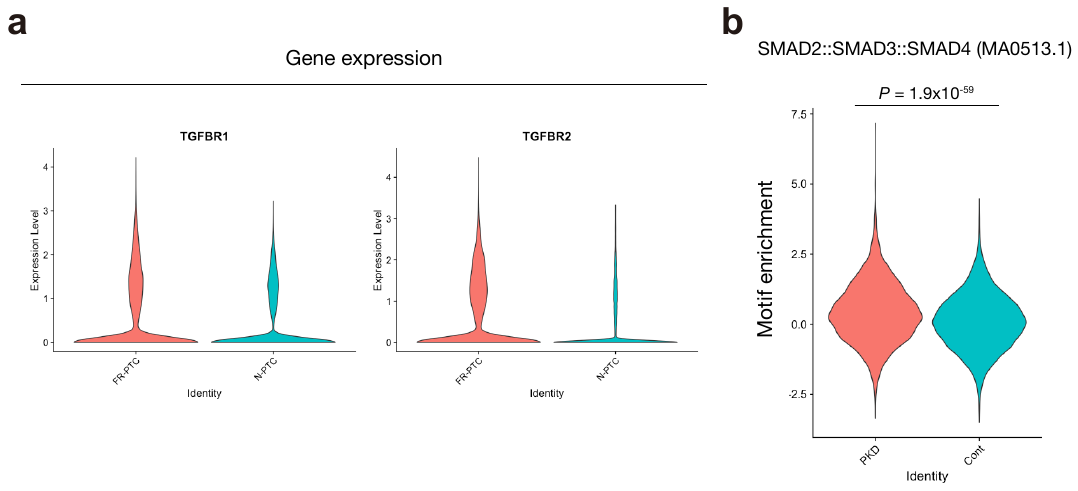

**Supplementary Figure 5. Up-regulation of TGFβ receptors in FR-PTC**

(**a**) Violin plot showing *TGFBR1* (left) or *TGFBR2* (right) gene expression in FR-PTC or N-PTC. (**b**) Violin plot showing SMAD2::SMAD3::SMAD4 binding motif (MA0513.1) enrichment score for PCT in ADPKD or control snATAC-seq dataset.

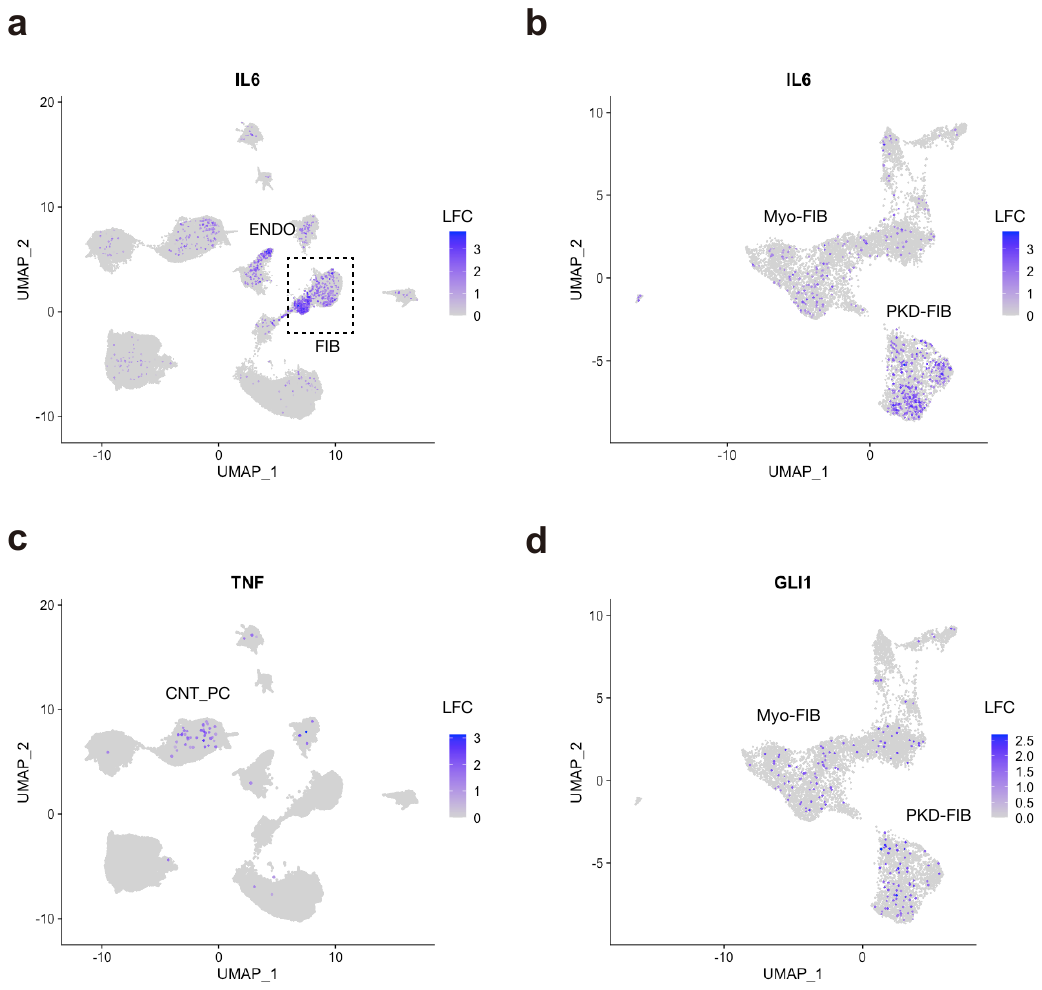

**Supplementary Figure 6. Unique molecular signature of ADPKD-specific fibroblast subtype**

(**a,b**) Umap plot displaying *IL6* expression in the whole dataset (**a**) or FIB subclusters (**b**). (**c**) Umap plot displaying *TNF* (TNFα) expression in the whole dataset. (**d**) Umap plot displaying *GLI1* expression in the FIB subclusters. The color scale for each plot represents a normalized log-fold-change (LFC).

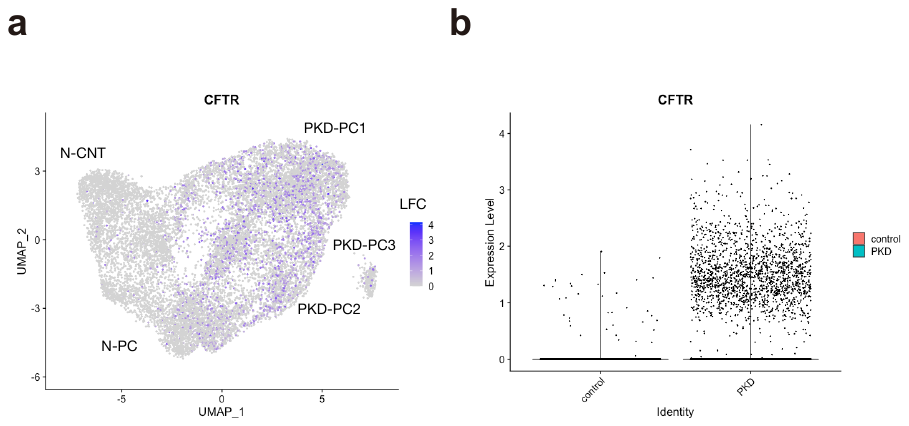

**Supplementary Figure 7. *CFTR* expressions in PC subclustering analysis**

(**a,b**) *CFTR* expression in CNT_PC subclustering analysis displayed with UMAP plot (**a**) or violin plot comparing ADPKD and control cells (**b**). The color scale for each plot represents a normalized log-fold-change (LFC).

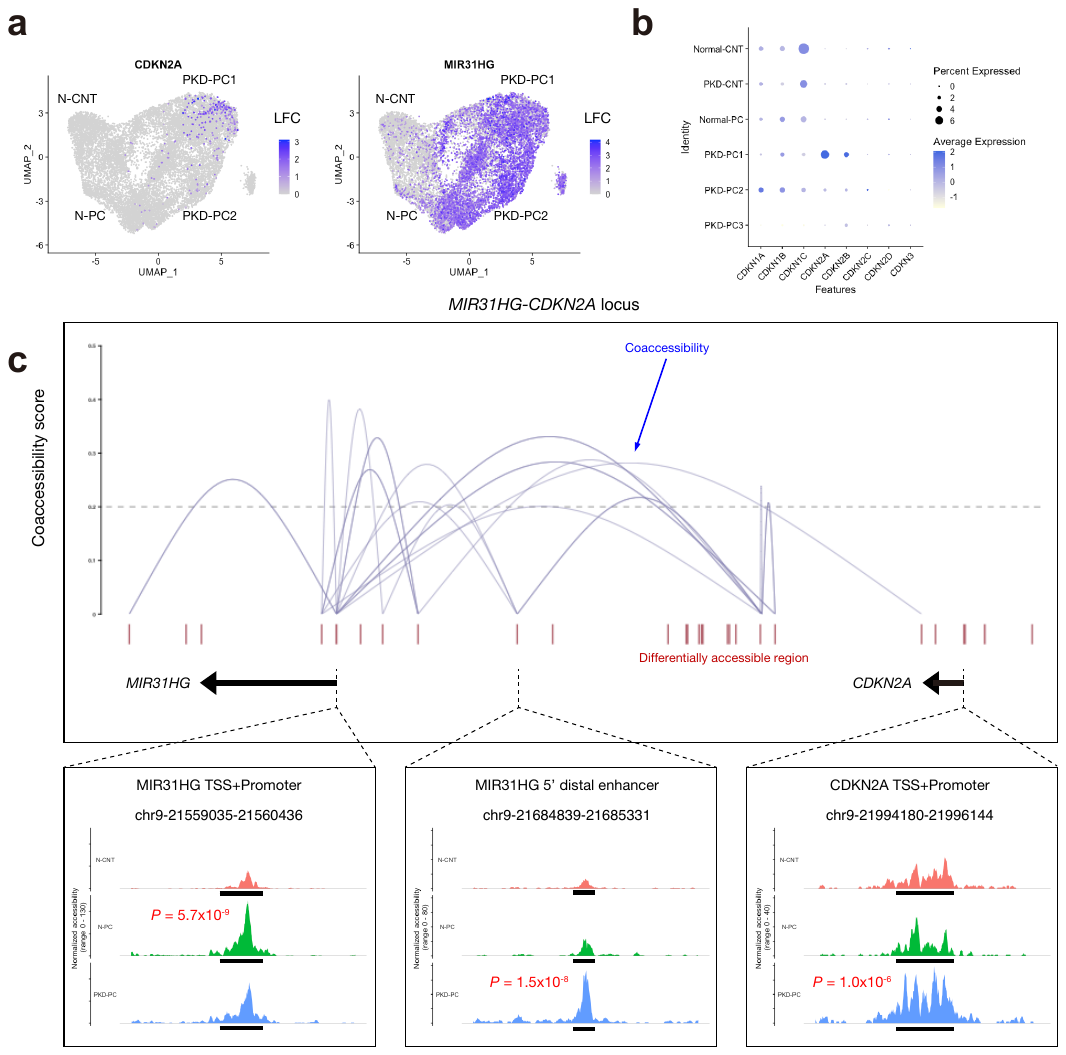

**Supplementary Figure 8. Multimodal approach identified anti-senescence gene enhancer activation in PC subpopulations in ADPKD**

(**a**) Umap plot displaying *CDKN2A* (left) or *MIR31HG* (right) gene expression in the snRNA-seq. The color scale for each plot represents a normalized log-fold-change (LFC). (**b**) Dot plots showing gene expression patterns of the cyclin-dependent kinase inhibitor genes enriched in each of PC subpopulations. (**c**) Cis-coaccessibility networks (CCAN, gray arcs) around the *MIR31HG* and *CDKN2A* gene in the ADPKD kidneys among accessible regions (red boxes) is shown (top). Fragment coverage (frequency of Tn5 insertion) around TSS (chr9:21559035-21560436) or 5' distal DAR (chr9:21684839-21685331) of *MIR31HG* and TSS of *CDKN2A* (chr9:21994180-21996144) are also shown (bottom, peak +/-3 Kb).

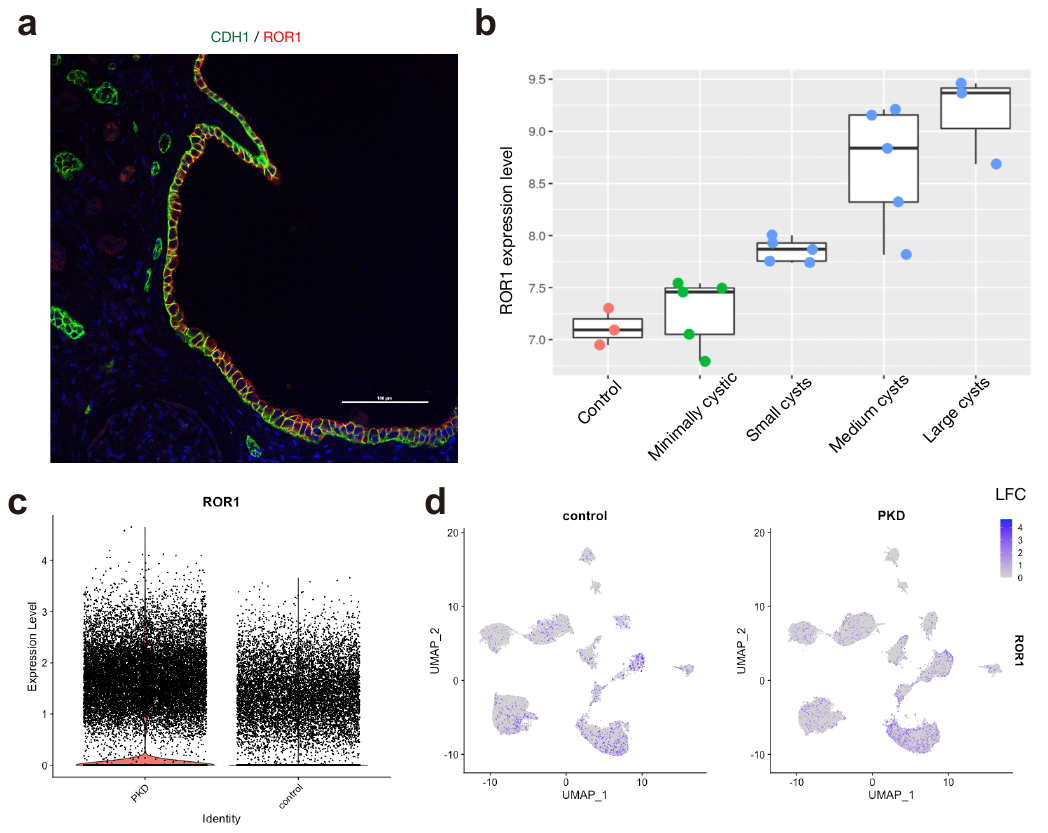

**Supplementary Figure 9. ROR1 expression in PC lineages of ADPKD kidneys**

(**a**) Representative immunohistochemical images of ROR1 (red) and CDH1 (green) in the ADPKD kidneys. Scale bar indicates 100 µm. (**b**) Correlation between *ROR1* expression and cyst size (GSE7869) (**c**) *ROR1* gene expression between ADPKD and control nuclei in all cell types. (**d**) Umap plot displaying *ROR1* gene expression in the control dataset (left) or ADPKD dataset (right). The color scale for each plot represents a normalized log-fold-change (LFC).

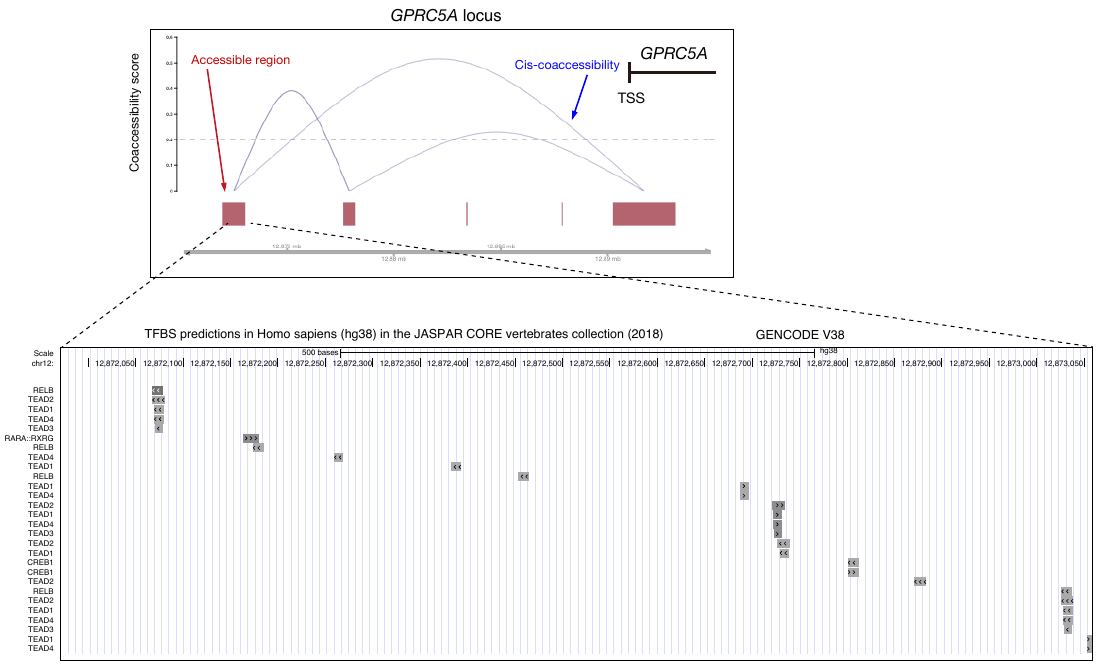

**Supplementary Figure 10. Enrichment of NF-κB, TEAD, CREB and retinoic acid receptor family transcription factor binding motifs on 5' distal region of *GPRC5A* gene**

The 5' distal region of *GPRC5A* gene (upper panel) has several binding motifs for cAMP responsive element binding protein 1 (CREB1) and retinoic acid receptors (RAR) as well as NF-κB (RELB) and TEAD family transcription factors (TEAD1-4), based on TFBS predictions in Homo sapiens (hg38) in the JASPAR CORE vertebrates collection (2018) shown on the UCSC genome browser (Enrichment score > 300, lower panel).

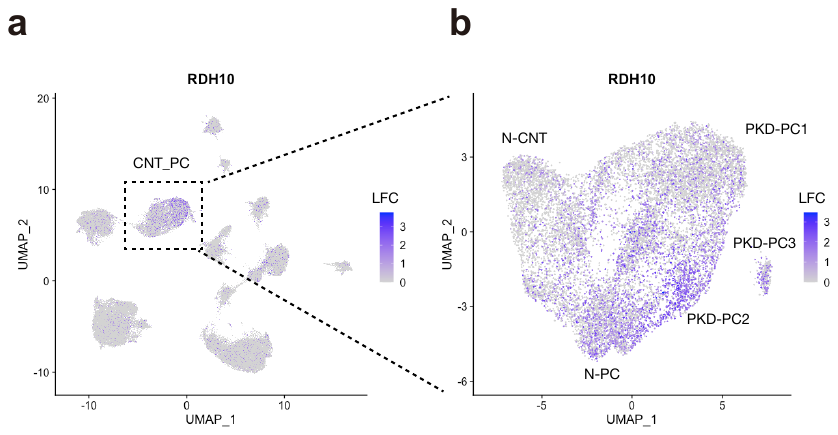

**Supplementary Figure 11. *RDH10* expression in principal cell lineages in ADPKD kidneys**

(**a,b**) Umap plot displaying *RDH10* expression in the whole dataset (**a**) or PC subclusters (**b**). The color scale for each plot represents a normalized log-fold-change (LFC).

| **Sample ID** | **Gender** | **Age** | **Pre-emptive transplant or chronic dialysis prior to transplant** | **Duration of sample preservation (day)** | **Kidney size** |
| --- | --- | --- | --- | --- | --- |
| PKD1 | Female | 56 | Dialysis | 810 | 1079 g |
| PKD2 | Female | 53 | Pre-emptive | 1207 | 803 g |
| PKD3 | Female | 58 | Pre-emptive | 216 | 1207 g |
| PKD4 | Male | 54 | Pre-emptive | 988 | 2061 g |
| PKD5 | Female | 61 | Pre-emptive | 535 | 1094 g |
| PKD6 | Male | 35 | Pre-emptive | 549 | 3170 g |
| PKD7 | Male | 42 | Pre-emptive | 629 | 2100 g |
| PKD8 | Male | 47 | Pre-emptive | 570 | 1531 g |

**Supplementary Table 1. Patient demographics and clinical information abstracted from the medical record**

Pre-emptive transplant or chronic dialysis prior to transplant, whether the patient was on maintenance dialysis or not (pre-emptive transplantation). Duration of sample preservation (day), duration (days) between sample collection and snRNA-seq library preparation.
collect the sample; Date of library prep, date of snRNA-seq

| snRNA-seq | Control | | ADPKD | |
| --- | --- | --- | --- | --- |
|  | Number | Frequency | Number | Frequency |
| PT+FR-PTC | 10523 | 25.9% | 12649 | 20.4% |
| PEC | 869 | 2.1% | 1634 | 2.6% |
| TAL1 | 13030 | 32.1% | 11253 | 18.1% |
| TAL2 | 852 | 2.1% | 1265 | 2.0% |
| DCT | 6225 | 15.3% | 4060 | 6.5% |
| CNT_PC | 3747 | 9.2% | 11143 | 18.0% |
| ICA | 1096 | 2.7% | 1900 | 3.1% |
| ICB | 1014 | 2.5% | 361 | 0.6% |
| PODO | 624 | 1.5% | 1585 | 2.6% |
| ENDO | 1693 | 4.2% | 3829 | 6.2% |
| FIB | 625 | 1.5% | 8244 | 13.3% |
| LEUK | 339 | 0.8% | 4064 | 6.5% |
| URO1 | 0 | 0.0% | 65 | 0.1% |
| URO2 | 0 | 0.0% | 21 | 0.0% |
| Total | 40637 | 100% | 62073 | 100% |

**Supplementary Table 2. The number or frequency of nuclei for each cell type quantitated in the whole filtered snRNA-seq dataset:** PT, proximal tubule; FR-PTC, failed-repair proximal tubular cells; PEC, parietal epithelial cells; TAL, thick ascending limb of Henle's loop; DCT, distal convoluted tubule; CNT_PC, connecting tubule and principle cells, ICA, Type A intercalated cells; ICB, Type B intercalated cells; PODO, podocyte; ENDO, endothelial cells; FIB, fibroblasts; LEUK, leukocytes; URO, uroepithelium.

| snATAC-seq | Control | | ADPKD | |
| --- | --- | --- | --- | --- |
|  | Number | Frequency | Number | Frequency |
| PCT | 9176 | 27.3% | 829 | 4.8% |
| PST | 3537 | 10.5% | 96 | 0.6% |
| FR-PTC | 1198 | 3.6% | 3119 | 18.0% |
| PEC_PODO | 746 | 2.2% | 292 | 1.7% |
| TAL | 9289 | 27.6% | 3406 | 19.6% |
| DCT | 3095 | 9.2% | 287 | 1.7% |
| CNT_PC | 3036 | 9.0% | 2765 | 15.9% |
| IC | 1509 | 4.5% | 615 | 3.5% |
| ENDO | 1480 | 4.4% | 1037 | 6.0% |
| FIB | 498 | 1.5% | 3241 | 18.7% |
| LEUK | 57 | 0.2% | 1678 | 9.7% |
| Total | 33621 | 100.0% | 17365 | 100.0% |

**Supplementary Table 3. The number or frequency of nuclei for each cell type quantitated in the whole filtered snATAC-seq dataset:** PCT, proximal convoluted tubule; PST, proximal straight tubule; FR-PTC, failed-repair proximal tubular cells; PEC-PODO, parietal epithelial cells and podocyte; TAL, thick ascending limb; DCT, distal convoluted tubule; CNT_PC, connecting tubule and principle cells, ICA, Type A and TypeB intercalated cells; ENDO, endothelial cells; FIB, fibroblasts; LEUK, leukocytes.

| **Donor** | **Total RNA reads** | **RNA reads per cell** | **Total ATAC reads** | **ATAC fragments per cell** |
| --- | --- | --- | --- | --- |
| Control_1 | 243,702,726 | 27,226 | 343,687,555 | 13,892 |
| Control _2 | 474,983,824 | 30,658 | 269,154,397 | 12,611 |
| Control _3 | 212,996,173 | 22,783 | 396,525,222 | 17,493 |
| Control _4 | 977,094,922 | 73,782 | 314,024,252 | 10,567 |
| Control _5 | 132,744,441 | 13,696 | 267,097,032 | 10,168 |
| ADPKD_1 | 297,480,919 | 24,616 | 324,886,233 | 9,755 |
| ADPKD_2 | 460,253,422 | 31,274 | 372,923,215 | 39,809 |
| ADPKD_3 | 370,314,010 | 31,171 | 374,516,045 | 26,190 |
| ADPKD_4 | 334,024,895 | 22,458 | 253,540,663 | 9,233 |
| ADPKD_5 | 303,934,085 | 28,312 | 282,728,864 | 11,359 |
| ADPKD_6 | 422,011,072 | 25,609 | 348,954,290 | 27,627 |
| ADPKD_7 | 307,102,696 | 93,429 | 518,627,214 | 34,513 |
| ADPKD_8 | 372,678,869 | 101,520 | 201,042,995 | 14,884 |

**Supplementary Table 4. The numbers of total reads and reads per nucleus in snRNA-seq and snATAC-seq data:** Total reads and reads per cell in snRNA-seq and snATAC-seq data were shown.

| **Quality Control for snRNA libraries** | | |
| --- | --- | --- |
| **Donor** | **Sequencing Saturation** | **Fraction reads with Valid Barcode** |
| Control_1 | 52.6 | 55.8 |
| Control _2 | 49.7 | 64.7 |
| Control _3 | 39.5 | 48.0 |
| Control _4 | 65.7 | 51.8 |
| Control _5 | 30.4 | 49.1 |
| ADPKD_1 | 45.4 | 38.4 |
| ADPKD_2 | 44.2 | 41.2 |
| ADPKD_3 | 46.6 | 40.1 |
| ADPKD_4 | 57.7 | 36 |
| ADPKD_5 | 34.1 | 37.9 |
| ADPKD_6 | 44.5 | 44.5 |
| ADPKD_7 | 73.8 | 25.9 |
| ADPKD_8 | 77.4 | 31.5 |

**Supplementary Table 5 – Quality control for snRNA-seq libraries:** The library complexity for the snRNA libraries was estimated with sequencing saturation for each donor. The fraction of read with a valid barcode in each donor.

| **Quality Control for snATAC libraries** | | |
| --- | --- | --- |
| **Donor** | **Sequencing Saturation** | **Fraction reads with Valid Barcdode** |
| Control_1 | 36.2 | 98.3 |
| Control _2 | 35.1 | 98.3 |
| Control _3 | 37.1 | 98.3 |
| Control _4 | 41.1 | 95.8 |
| Control _5 | 37.3 | 95.8 |
| ADPKD_1 | 25.6 | 95.4 |
| ADPKD_2 | 40.5 | 97.1 |
| ADPKD_3 | 31.9 | 97.7 |
| ADPKD_4 | 17.4 | 95.8 |
| ADPKD_5 | 32.7 | 95.8 |
| ADPKD_6 | 27.1 | 96.9 |
| ADPKD_7 | 49 | 97.2 |
| ADPKD_8 | 28.4 | 85.8 |

**Supplementary Table 6 – Quality control for snATAC-seq libraries:** The sequencing saturation and the fraction of reads with a valid barcode in each donor were shown.

| **Oligonucleotides cloned for sgRNA expression** | | |
| --- | --- | --- |
| Enhancer #1  (chr12:12872917) | Forward | CACCGACCTTCAGGGTCGCCTAACT |
|  | Reverse | AAACAGTTAGGCGACCCTGAAGGTC |
| Enhancer #2  (chr12:12872506) | Forward | CACCGTCTACCGGTTTATGTGTATA |
|  | Reverse | AAACTATACACATAAACCGGTAGAC |
| Promoter #1  (chr12:12891514) | Forward | CACCGTAAAGGCGGCCCTCGCCGGA |
|  | Reverse | AAACTCCGGCGAGGGCCGCCTTTAC |
| Promoter #2  (chr12:12891297) | Forward | CACCGTCGGAGGAGTCCGATGCGCT |
|  | Reverse | AAACAGCGCATCGGACTCCTCCGAC |
| Non-targeting #1 | Forward | CACCGGTAAGCGCGTGAGTCGAA |
|  | Reverse | AAACTTCGACTCACGCGCTTACC |
| Non-targeting #2 | Forward | CACCGAGGCGAGGTAAGACGCGG |
|  | Reverse | AAACCCGCGTCTTACCTCGCCTC |

**Supplementary Table 7 – Oligonucleotides cloned for sgRNA expression**
